# Estimation of Genetic Admixture Proportions via Haplotypes

**DOI:** 10.1101/2023.09.22.559067

**Authors:** Seyoon Ko, Eric M. Sobel, Hua Zhou, Kenneth Lange

## Abstract

Estimation of ethnic admixture is essential for creating personal genealogies, studying human history, and conducting genome-wide association studies (GWAS). Three methods exist for estimating admixture coefficients. The frequentist approach directly maximizes the binomial loglikelihood. The Bayesian approach adds a reasonable prior and samples the posterior distribution. Finally, the nonparametric approach decomposes the genotype matrix algebraically. Each approach scales successfully to data sets with a million individuals and a million single nucleotide polymorphisms (SNPs). Despite their variety, all current approaches assume independence between SNPs. To achieve independence requires performing LD (linkage disequilibrium) filtering before analysis. Unfortunately, this tactic loses valuable information and usually retains many SNPs still in LD. The present paper explores the option of explicitly incorporating haplotypes in ancestry estimation. Our program, HaploADMIXTURE, operates on adjacent SNP pairs and jointly estimates their haplotype frequencies along with admixture coefficients. This more complex strategy takes advantage of the rich information available in haplotypes and ultimately yields better admixture estimates and better clustering of real populations in curated data sets.

## 1. Introduction

Estimation of genetic admixture is key to reconstructing personal genealogies and understanding population histories^1^. Adjusting for genetic ancestry is also a necessary prelude to genome-wide association studies (GWAS) for medical and anthropological traits ^2^. Failure to account for ancestry can lead to false positives due to population stratification ^3,4,5^. In these analyses, admixture coefficients serve as covariates adjusting for ancestry. Because admixture coefficients represent the proportions of a person’s ancestry derived from different founding populations, they are more interpretable than principal components (PCs).

Admixture coefficients can be estimated simultaneously with allele frequencies in known or latent populations. ADMIXTURE^6^ is the most widely-used likelihood-based software. It directly maximizes the binomial likelihood of the admixture coefficients and allele frequencies via alternating sequential quadratic programming^7^. Our recent Julia version, OpenADMIXTURE^8^, incorporates time-saving software enhancements and AIM (ancestry informative markers) preselection via sparse *K*-means clustering ^9^. STRUCTURE^10^ and its extensions fastStructure^11^ and TeraStructure^12^ rely on Bayesian inference. SCOPE^13^ replaces the genotype matrix by a low-rank matrix, which is delivered by alternating least squares^14^ and randomized linear algebra. Each of the recent versions of these programs – OpenADMIXTURE, TeraStructure, and SCOPE – scales to biobank-size data sets of a million people and a million single nucleotide polymorphisms (SNPs).

A regrettable limitation of most of these programs is their assumption of independence for the alleles present at neighboring SNPs. To avoid this patently false assumption, SNPs are filtered to remove SNPs in linkage disequilibrium (LD). Filtering must reach a balance between LD elimination and the loss of valuable AIMs. Unfortunately, the LD-aware program fineSTRUCTURE^15^ scales poorly on large data sets^13^. Yet the evidence is clear that ancestry informative haplotypes pack more information than ancestry informative SNPs^16,17,18^.

The current paper demonstrates the value of haplotypes in admixture estimation and population clustering. Given the combinatorial and computational complexities encountered, we consider only haplotypes formed from adjacent SNP pairs. Even with this limitation, haplotype models offer substantial improvements in estimation and clustering in simulated and real data sets on well-separated ancestral populations. Our new program, HaploADMIXTURE, builds on the high-performance computing (HPC) techniques pioneered in OpenADMIXTURE^8^. By leveraging the parallel processing capabilities of graphics processing units (GPUs), HaploADMIXTURE is able to run in reasonable time. In practice only a minority of haplotypes are informative. To select ancestry informative haplotypes, we exploit unsupervised sparse *K*-means clustering via feature ranking ^8,9^. This generic method, denoted by the acronym SKFR, selects the informative features (haplotypes) driving cluster formation. Our experience suggests that the SKFR-HaploADMIXTURE pipeline delivers the best admixture results currently available with reasonable computing times.

## 2. Methods

### 2.1 Admixture Likelihoods

Consider a sample of *I* unrelated individuals, *B* haplotype blocks, and *S* SNPs per block. For our purposes *S* equals 1 or 2. Let ***x***_*ib*_ denote the length-*S* genotype vector for haplotype block *b* of individual *i*. Each genotype of *i* counts the number of *i*’s reference alleles present and is coded as a number from the set {0, 1, 2}. Haplotypes are coded as sequences of 0’s and 1’s, and every ***x***_*ib*_ = ***h***_*ib*1_ + ***h***_*ib*2_ equals a sum of a maternal and paternal haplotypes. The blocks are taken to be contiguous, non-overlapping, and exhaustive. Haplotypes may be chosen through feature selection as discussed in Section 2.4. Let *p*_*kbh*_ *>* 0 be the frequency of haplotype ***h*** of haplotype block *b* in population *k*, and let *q*_*ki*_ *>* 0 denote the fraction of *i*’s genome coming from population *k*, where 1 ≤ *k* ≤ *K*. The loglikelihood of the sample under a binomial distribution and independence of haplotype blocks is

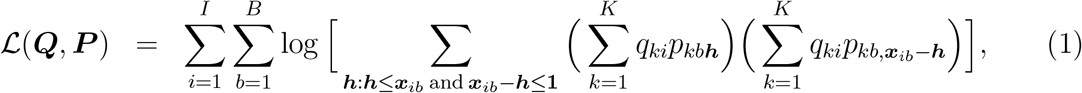

where ***x***_*ib*_ is the sum of the maternal haplotype **0** ≤ ***h*** ≤ **1** and the paternal haplotype ***x***_*ib*_ − ***h***. There are 2^*S*^ possible haplotypes per block, and the constraints 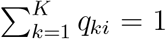 and Σ_***h***_ *p*_*kb**h***_ = 1 hold for each *i* and combination (*k, b*). The loglikelihood (1) simplifies by symmetry if any entry of ***x***_*ib*_ equals 1 (a heterozygous SNP). Because maternal and paternal haplotypes are interchangeable, the number of summands can be halved if the remaining sum of products is doubled. When *S* = 1 and *i* and *b* are fixed, the heterozygous genotype 1 has probability 2(Σ_*k*_ *q*_*ki*_*p*_*kb*0_)(Σ_*k*_ *q*_*ki*_*p*_*kb*1_), which the log function splits into a sum of logarithms. In fact, this simplification replicates the binomial likelihood employed in ADMIXTURE and STRUCTURE. When *S* = 2, the doubly heterozygous genotype has the probability 2[Σ_*k*_ *q*_*ki*_*p*_*kb*(00)_] [Σ_*k*_ *q*_*ki*_*p*_*kb*(11)_] +2[Σ_*k*_ *q*_*ki*_*p*_*kb*(01)_] [Σ_*k*_ *q*_*ki*_*p*_*kb*(10)_], which no longer splits under the log function. In addition, there are cases where one of the genotypes is observed, but the other is missing. For example, if the first genotype is heterozygous and the other is missing, the probability equals [Σ_*k*_ *q*_*ki*_*p*_*kb*(00)_ +Σ_*k*_*q*_*ki*_*p*_*kb*(01)_] [Σ_*k*_ *q*_*ki*_*p*_*kb*(10)_ + Σ_*k*_ *q*_*ki*_*p*_*kb*(11)_], and the loglikelihood (1) should be adjusted accordingly. Nonetheless, as described in the next subsection, the whole loglikelihood is still amenable to maximization.

### 2.2 Maximum Likelihood Estimation

Estimation in HaploADMIXTURE and OpenADMIXTURE are similar. The optimization machinary in both programs alternate estimation of the per-population haplotype frequencies *p*_*kb**h***_ and the per-individual admixture coefficients *q*_*ki*_. To allow easy parallelization with graphics processing units (GPUs), we invoke the minorization-maximization (MM) principle^19,20^ to split sums appearing in the arguments to the logarithms of the haplotype loglikelihood (1). The operative inequality

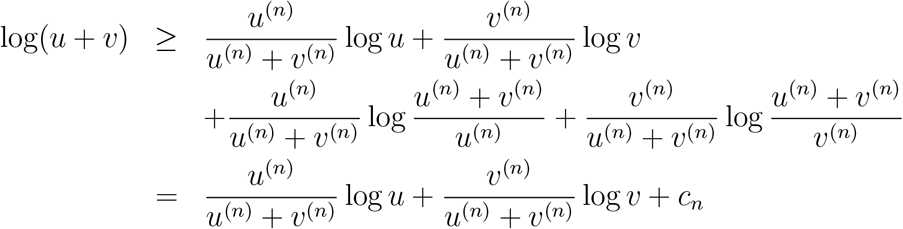

reduces to an equality when *u* = *u*^(*n*)^ and *v* = *v*^(*n*)^. Here the irrelevant constant *c*_*n*_ depends only on the current values *u*^(*n*)^ and *v*^(*n*)^ of *u* and *v*. The function 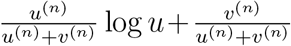 becomes a surrogate for the function log(*u* + *v*) it replaces. For example, when *S* = 2 and *i* presents a doubly heterozygous genotype, we take *u* = 2 [Σ_*k*_ *q*_*ki*_*p*_*kb*(00)_] [Σ_*k*_ *q*_*ki*_*p*_*kb*(11)_] and *v* = 2 [Σ_*k*_ *q*_*ki*_*p*_*kb*(01)_] [Σ_*k*_ *q*_*ki*_*p*_*kb*(10)_] . Most genotype probabilities (all homozygous and singly heterozygous genotypes) reduce to a single product where log splitting is unnecessary. For haplotypes involving more than two SNPs, phase combinations become more complex, code is harder to write, and computation slows. For these reasons we venture no further than two-SNP haplotypes. Maximization of the surrogate function created by minorization enjoys the ascent property of steadily increasing the loglikelihood. The ascent property is the essence of the MM (minorization-maximization) principle^19,20^.

Minorization creates a surrogate function

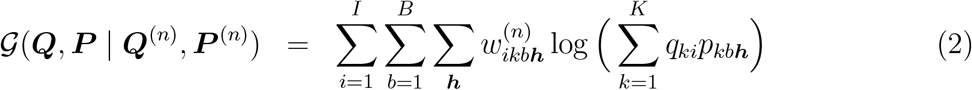

involving nonnegative weights 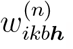, many of which are 0 because they correspond to haplotypes incompatible with observed genotypes. Except for revising the weights 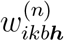 at each iteration *n*, the surrogate loglikelihood (2) is simpler to deal with than the actual loglikelihood. Updating the admixture matrix ***Q*** = (*q*_*ki*_) can be done simultaneously over columns (individuals *i*). Updating the haplotype frequency tensor ***P*** = (*p*_*kb**h***_) can be done simultaneously over its middle columns (blocks *b*). Each such maximization must respect the nonnegativity constraints on the proportions and their sum to 1 constraints. Very simple multinomial updates of the *p*_*kb**h***_ can be achieved by splitting the argument 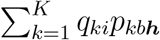 of the log function, but this second minorization slows convergence dramatically.

The parallel updates of ***P*** and ***Q*** are structured around functions of the form

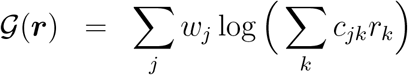

subject to nonnegativity and sum to 1 constraints. The method of recursive quadratic programming involves replacing G(***r***) by its local quadratic approximation

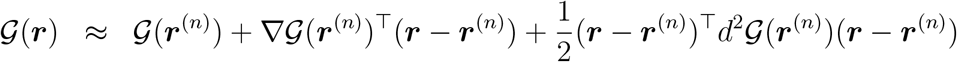

and maximizing this approximation subject to the constraints. The required gradient and Hessian are

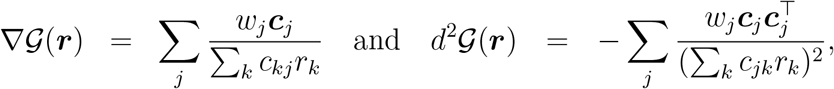

where 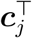 is the *j*th row of the matrix ***C*** of nonnegative constants *c*_*jk*_.

Computation of the gradients and Hessians of 𝒢(***Q, P*** | ***Q***^(*n*)^, ***P*** ^(*n*)^) with respect to ***Q*** has time complexity *O*(2^*S*^*IBK*^2^) and space complexity *O*(*IK*^2^). Computation of the gradients and Hessians with respect to ***P*** again has time complexity *O*(2^*S*^*IBK*^2^) but now space complexity *O*(2^*S*^*BK*^2^). The quadratic programming cost of updating ***Q*** breaks down into *I* quadratic programs of size *K* with a single equality constraint. The cost of solving one of these quadratic programs is *O*(*K*^3^). The quadratic programming cost of updating ***P*** breaks down into *B* quadratic programs of size 2^*S*^*K* with an equality constraint for each population *k*. The cost of solving one of these quadratic programs is *O*[(2^*S*^*K* + *K*)^3^]. In practice, when *S* = 2, the time needed for solving the quadratic programs for ***Q*** is negligible compared to the time needed for computing gradients and Hessians. In contrast, the time needed for solv-ing the quadratic programs for ***P*** is comparable to the time needed for computing gradients and Hessians.

Our Julia implementation of HaploADMIXTURE allows users to invoke Nvidia graphics processing units (GPUs) to accelerate the evaluation of gradients and Hessians and to solve the various quadratic programs. Convergence criteria can be set by the user. The default setting for overall convergence mandates that the relative change in loglikelihoods falls below 10^−7^.

### 2.3 Selection of *K*

We employ two devices to select the number of ancestral populations *K*. First, the cross-validation method introduced in ADMIXTURE^21^ partitions the sample individuals into *v* folds. Each of the folds is held out as a validation set, and the model is fit on the remaining training individuals. Fitting on a training set is fast because it warm starts parameter values from the estimates garnered under the full data set. Given the haplotype frequencies ***P***_*train*_ estimated on the training set, we estimate the admixture fractions ***Q***_*test*_ on the validation set. This fitting step is also fast because it qualifies as a straightforward convex supervised learning problem. Given ***P***_*train*_ and ***Q***_*test*_, we predict the genotype matrix of the validation individuals. The deviance residual under a binomial model yields the prediction error

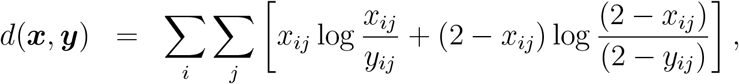

where ***x*** is *I* × *SB* true genotype matrix, and ***y*** is the predicted genotype matrix. This error is then averaged across the different folds. We choose the most parsimonious model whose prediction error is no more than one standard error above the error of the best model (one standard error rule).

The second device for selecting *K* is the Akaike information criterion (AIC)^22^. In the current setting

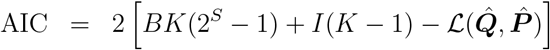

The term *BK*(2^*S*^ − 1) + *I*(*K* − 1) counts the number of free parameters in the model with *K* ancestral populations. The loglikelihood is evaluated at the maximum likelihood estimates given *K*. We fit the model for several different values of *K* and choose the *K* with the lowest value of AIC. The virtue of AIC is that it requires less computation than full cross-validation.

### 2.4 Sparse *K*-Means with Feature Ranking for Haplotypes

To select AIMs, sparse *K*-means with feature ranking (SKFR)^8,9^ has proved ideal. SKFR ranks and selects a predetermined number of features based on their importance in *K*-means clustering. HaploADMIXTURE requires input blocks of SNPs rather than individual SNPs. The center for cluster *j* is a vector ***c***_*j*_ = (*c*_*jg*_). The loss in *K*-means clustering is 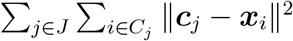, where *C*_*j*_ denotes the set of individuals belonging to cluster *j*, and each raw feature vector ***x***_*i*_ = ***h***_*m*_ + ***h***_*p*_ is a sum of unknown haplotypes. (If everyone is haplotyped, then SKFR should operate on haplotypes.) In practice, the ***x***_*i*_ are standardized to have a mean of 0 for each feature across the entire sample. A missing genotype *x*_*ig*_ in ***x***_*i*_ is imputed by *c*_*jg*_ when *i* is assigned to cluster *j* ^23^. To model haplotypes, the feature vector ***x***_*i*_ is broken into vector blocks ***x***_*ib*_. Lloyd’s algorithm^24^ alternates updating cluster centers and reassigning feature vectors to clusters. At each iteration of Lloyd’s algorithm, the *S* blocks giving the largest reduction in the loss are selected based on the decomposition ∥***c***_*j*_ − ***x***_*i*_∥^2^= Σ_*b*_ ∥***c***_*jb*_ − ***x***_*ib*_∥^2^. The mean for a selected block is cluster-specific. The mean for a non-selected block is taken to be **0**. Lloyd’s algorithm converges when the cluster centers and ancestry informative blocks stabilize.

### 2.5 Computational Tactics

Most of the computational tactics introduced in OpenADMIXTURE carry over to HaploADMIXTURE. For instance, HaploADMIXTURE significantly reduces memory demands by directly converting the bit genotypes stored in PLINK BED format ^25^ into numbers through the OpenMendel^26^ package SnpArrays^27^. Multithreading is employed throughout HaploADMIXTURE. Multithreading not only promotes parallelism, but also reduces memory usage by tiling the computation of gradients and Hessians. CUDA GPU kernels are implemented for EM updates and computing gradients and Hessians. When running SKFR for multiple sparsity levels *S*, we start with the highest level of *S* and warm start Lloyd’s algorithm at the current level by its converged value at the previous higher level. We refer the readers to Ko et al. ^8^ for further details.

### 2.6 Performance Evaluation

#### 2.6.1 Permutation Matching of Clusters

A promising similarity metric proposed by Behr et al. ^28^ is effective in matching clusters defined by two admixture matrices ***Q***^1^ and ***Q***^2^. This metric faithfully matches similar clusters and is invariant when cluster labels are permuted. The metric quantifies the similarity between cluster *m* in ***Q***^1^ and cluster *n* in ***Q***^2^ as

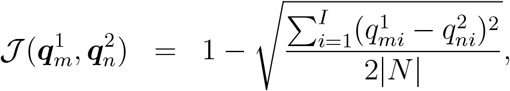

where *N* is the set of indices *i* for which 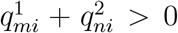, and |*N* | is the cardinality of *N* . To match the clusters delivered by two algorithms, we solve the assignment problem that maximizes the criterion 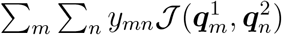, subject to the constraints *y*_*mn*_ ∈ {0, 1} and 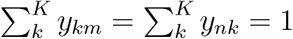, where *K* is the number of clusters. In practice, we relax the domain of *y*_*mn*_ to the unit interval and solve the simplified problem using linear programming via JuMP^29^, a mathematical optimization package in Julia.

#### 2.6.2 Silhouette Coefficient

The silhouette index *s*_*i*_ is a measure of how similar object *i* is to its own cluster (cohesion) compared to other clusters (separation)^30^. If *i* belongs to cluster *C*_*k*_, then the index *s*_*i*_ reflects the two averages

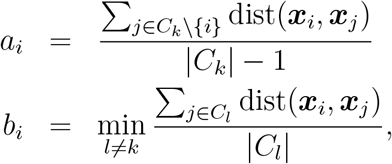

where *a*_*i*_ is the average distance of sample *i* from the other points in *C*_*k*_, and *b*_*i*_ is the minimum average distance of sample *i* from the other clusters. Given these values we define

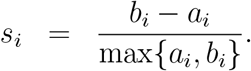

Note that *s*_*i*_ ranges from −1 to 1; the higher *s*_*i*_ is, the better separated the clusters are. Thus, the average silhouette value serves as a sensitive measure of clustering quality.

#### 2.6.3 Visualization

Stacked bar plots allow easy visualization of estimated admixture proportions when clusters are matched consistently across computer runs. Matching is accomplished by hierarchical clustering with complete linkage based on the HaploADMIXTURE ***Q*** estimates. Hierarchical clustering determines the order of samples within a population. One can also apply hierarchical clustering to the set of populations and to the set of continents. In the former case clustering operates on cluster centers and in the latter case on averages of cluster centers.

### 2.7 Real Data Sets

To evaluate its performance, we applied HaploADMIXTURE to three different real-world data sets: the 1000 Genomes Project (TGP)^31,32^, the Human Genome Diversity Project (HGDP)^33,34^, and the Human Origins (HO) ^35^ project. (We adhered to compliance agreements in each case.) The TGP data set includes genotypes from the 2012-01-31 Omni Platform after filtering to exclude related individuals, individuals with less than a 95% genotyping success rate, and variants with minor allele frequency (MAF) less tha 1%. The filtered data set contains 1718 unrelated individuals and 1,854,622 SNPs. The self-reported ancestry labels range over 26 different populations grouped into continental populations of African (AFR), Admixed American (AMR), East Asian (EAS), European (EUR), and South Asian (SAS) descent. The HGDP data set contains 940 individuals across 32 self-reported populations and 642,950 SNPs after filtering by the same criteria applied to the TGP data. The self-reported population labels are further grouped into seven continental labels: Europe, Middle East, Central South Asia, East Asia, Africa, America, and Oceania. The HO data set includes 1931 individuals across 163 populations and 385,088 SNPs. Here, filtering excludes individuals with less than 99% genotyping success rate and SNPs with MAF less than 5%. No continental population labels are provided for HO. We illustrate our results on the TGP data set. The corresponding results on HGDP and HO appear in the Supplementary Materials.

### 2.8 Simulations

A common approach for simulating genetic admixture with *S* = 1 is the Pritchard-Stephens-Donnelly (PSD) model^10^, with allele frequencies sampled from the Balding-Nicolas model ^36^ following a beta distribution:

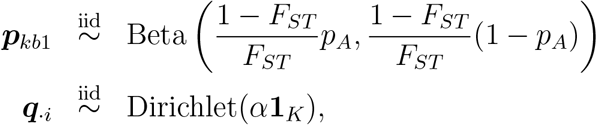

where *p*_*A*_ is the allele frequency and *F*_*ST*_ is the fixation index. Chiu et al. ^13^ introduced an extra level of Dirichlet sampling to simulate populations gathered around regional centers. This is accomplished by first sampling *T* regional centers ***s***_*t*_ from the Dirichlet(*α***1**_*K*_) distri-bution. Then for each regional center, *I/T* of the admixture vectors ***q***_·*i*_ are sampled around the center **s**_*t*_, with a high value of the parameter *γ* = 50. To model pairs of SNPs (*S* = 2), we simulated data according to a variant of the model for *S* = 1 with the beta distributions for allele frequencies replaced by the Dirichlet distributions for haplotype frequencies. Specifically, we generated

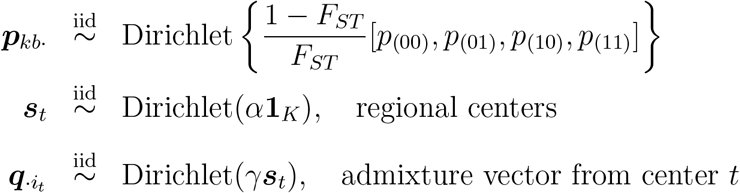

given the initial haplotype frequencies [*p*_(00)_, *p*_(01)_, *p*_(10)_, *p*_(11)_] and fixation indexes *F*_*ST*_ . To specify these underlying parameters, we randomly sampled a block of SNPs of length *S* from chromosome 1 of the TGP data set and captured their haplotype frequencies and the harmonic means of their estimated fixation indices. For admixture proportions *q*_*ki*_, different values of the parameter *α* determine the dispersion of the regions. A low value of *α* tends to generate more homogeneous populations, as shown in Figure S1. Finally, the haplotypes were sampled from

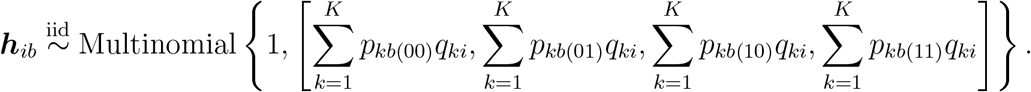

If any haplotype frequency *p*_*kb**h***_ fell outside the interval [0.005, 0.995], then we clamped it to the closest endpoint 0.005 or 0.995 and renormalized the frequencies for block *b* of population *k* to sum to 1.

## 3 Results

### 3.1 Simulation Studies

We simulated independent data sets with different numbers of SNPs and different values of the concentration parameter *α* ∈ {0.1, 0.05, 0.02} as described in Section 2.8. Tables 1, S3, and S2 display root-mean-square errors (RMSE) for 0,000, 100,000, and 1,000,000 simulated SNPs. RMSE is estimated by

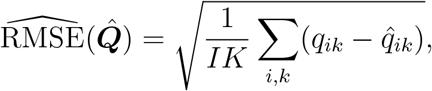

where ***Q*** are the true values and 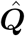 are the estimates. In all cases, both HaploADMIXTURE and OpenADMIXTURE perform as well as SCOPE and TeraStructure. HaploADMIXTURE shows better accuracy than OpenADMIXTURE in most cases. OpenADMIXTURE outperforms for 1,000,000 SNPs under relatively heterogeneous populations (*α* = 0.1). When AIMs selected by sparse *K*-means are input, HaploADMIXTURE shows less performance degradation than OpenADMIXTURE, sometimes even improving the results found with all of the SNPs. For all of the cases evaluated, both AIC and cross-validation correctly select *K* = 5 based on the one standard error rule.

**Table 1.** Root-mean-square errors of the estimated admixture proportions on the simulated data sets. Root-mean-square error checks the accuracy of the estimated admixture coefficients; the lower, the better. Five populations were used for the simulation, with 1000 individuals and 10,000 SNPs for various values of *α*. Each value is averaged over five simulation runs. The best value for each *α* is in italics.

| AIMs |  | HaploADMIXTURE | OpenADMIXTURE | SCOPE | TeraStructure |
| --- | --- | --- | --- | --- | --- |
| 10,000 SNPs, $\alpha = 0.1$ | | | | | |
|  | 10,000 | *0.0201* | 0.0300 | 0.0534 | 0.0630 |
|  | 2000 | 0.0290 | 0.0362 |  |  |
|  | 1500 | 0.0315 | 0.0383 |  |  |
|  | 1000 | 0.0357 | 0.0421 |  |  |
|  | 500 | 0.0446 | 0.0498 |  |  |
| 10,000 SNPs, $\alpha = 0.05$ | | | | | |
|  | 10,000 | *0.0194* | 0.0323 | 0.0582 | 0.0636 |
|  | 2000 | 0.0271 | 0.0371 |  |  |
|  | 1500 | 0.0293 | 0.0389 |  |  |
|  | 1000 | 0.0330 | 0.0422 |  |  |
|  | 500 | 0.0411 | 0.0488 |  |  |
| 10,000 SNPs, $\alpha = 0.02$ | | | | | |
|  | 10,000 | *0.0183* | 0.0327 | 0.0602 | 0.0539 |
|  | 2000 | 0.0255 | 0.0363 |  |  |
|  | 1500 | 0.0275 | 0.0378 |  |  |
|  | 1000 | 0.0307 | 0.0402 |  |  |
|  | 500 | 0.0376 | 0.0459 |  |  |

### 3.2 Real-world Data Sets

#### 3.2.1 Selection of *K*

To assess the performance of HaploADMIXTURE, we computed AIC values and performed cross-validation to select the best *K* for the real-world data sets. For TGP, both AIC and cross-validation select *K* = 7, while TeraStructure selects *K* = 8. For HGDP, AIC selects *K* = 7, but cross-validation and TeraStructure select *K* = 10. For HO, AIC selects *K* = 12, cross-validation selects *K* = 10, and TeraStructure selects *K* = 14. On balance, we prefer AIC because of its computational efficiency and parsimony. This preference is bolstered by the notable differences observed in the graphs between TeraStructure and OpenADMIXTURE covered in Section 3.2.2.

#### 3.2.2 Visualization

Figures 1, S2, and S3 illustrate the admixture proportions inferred from the TGP, HGDP, and HO data sets by HaploADMIXTURE, OpenADMIXTURE, SCOPE, and TeraStructure. The general structure seems similar across the programs, with some differences. For example, TeraStructure tends to rely on a single European (EUR) population in TGP, while the other programs tend to rely on two. Section 3.2.4 summarizes the ability of the programs to separate self-identified populations. Previous publications of Chiu et al. ^13^ and Ko et al. ^8^ incorrectly match individuals to populations because of a data reading error. Figure S2 fixes this error and clearly separates the different continental populations.

**Figure 1.**
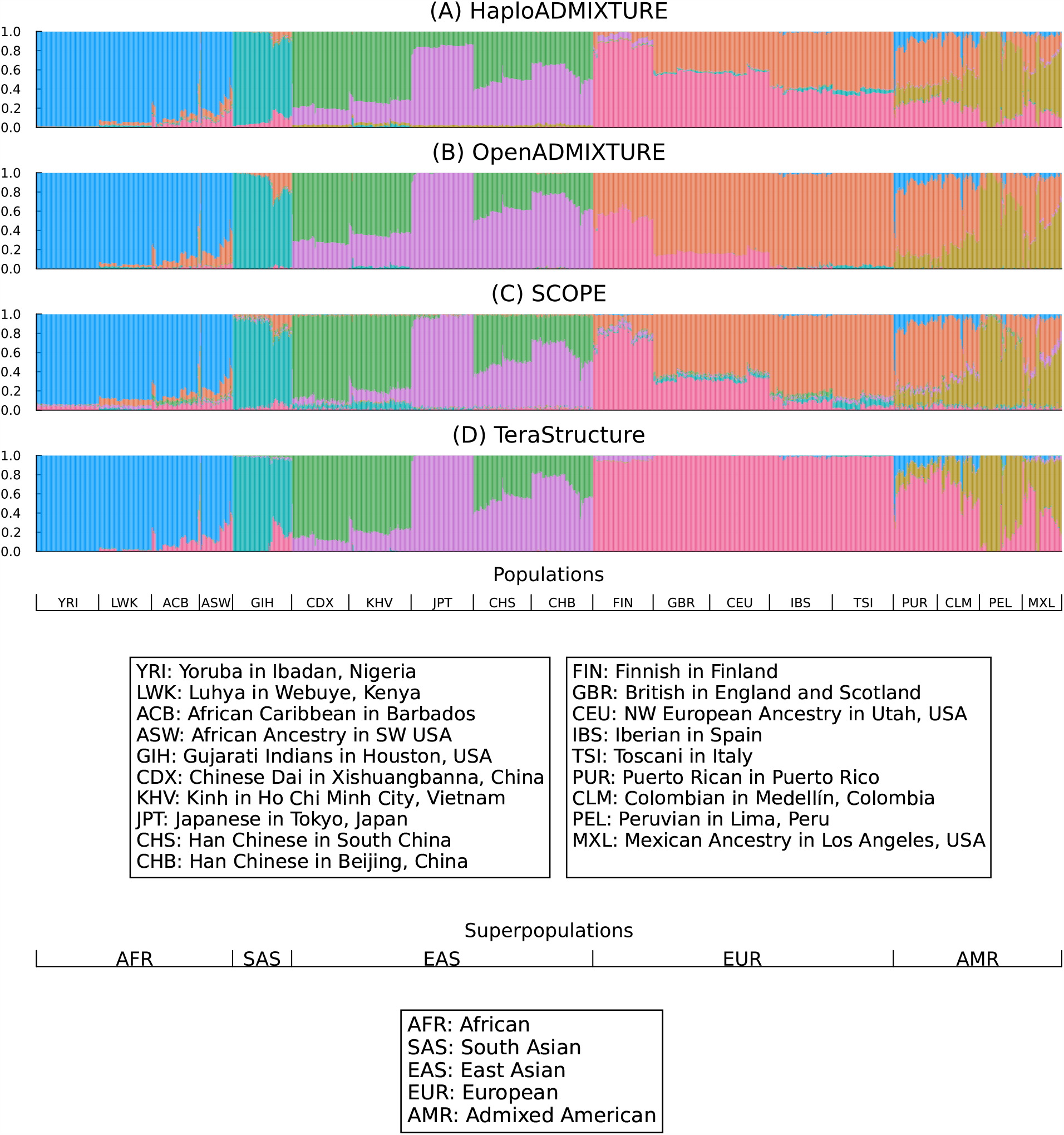
Ancestry estimation of TGP data samples. (a) Using HaploADMIXTURE with all SNPs, (b) OpenADMIXTURE with all SNPs, (c) SCOPE, and (d) TeraStructure. The results are presented in stacked bar plots, where the y-axis indicates the proportion of total ancestry. The x-axis shows all samples arranged by population labels.

Figures 2, S4, and S6 show the structures inferred by HaploADMIXTURE operating on AIMs chosen through sparse *K*-means clustering. Figures 3, S5, and S7 display the structures inferred by OpenADMIXTURE in the same circumstances. Evidently, HaploADMIXTURE faithfully reproduces the general structure with fewer AIMs than OpenADMIXTURE. In particular, OpenADMIXTURE fails to distinguish European populations from Middle-Eastern populations. Figures 4, S8, and S9 display the structure inferred by HaploADMIXTURE for different numbers of populations *K* as discussed in Section 3.2.1.

**Figure 2.**
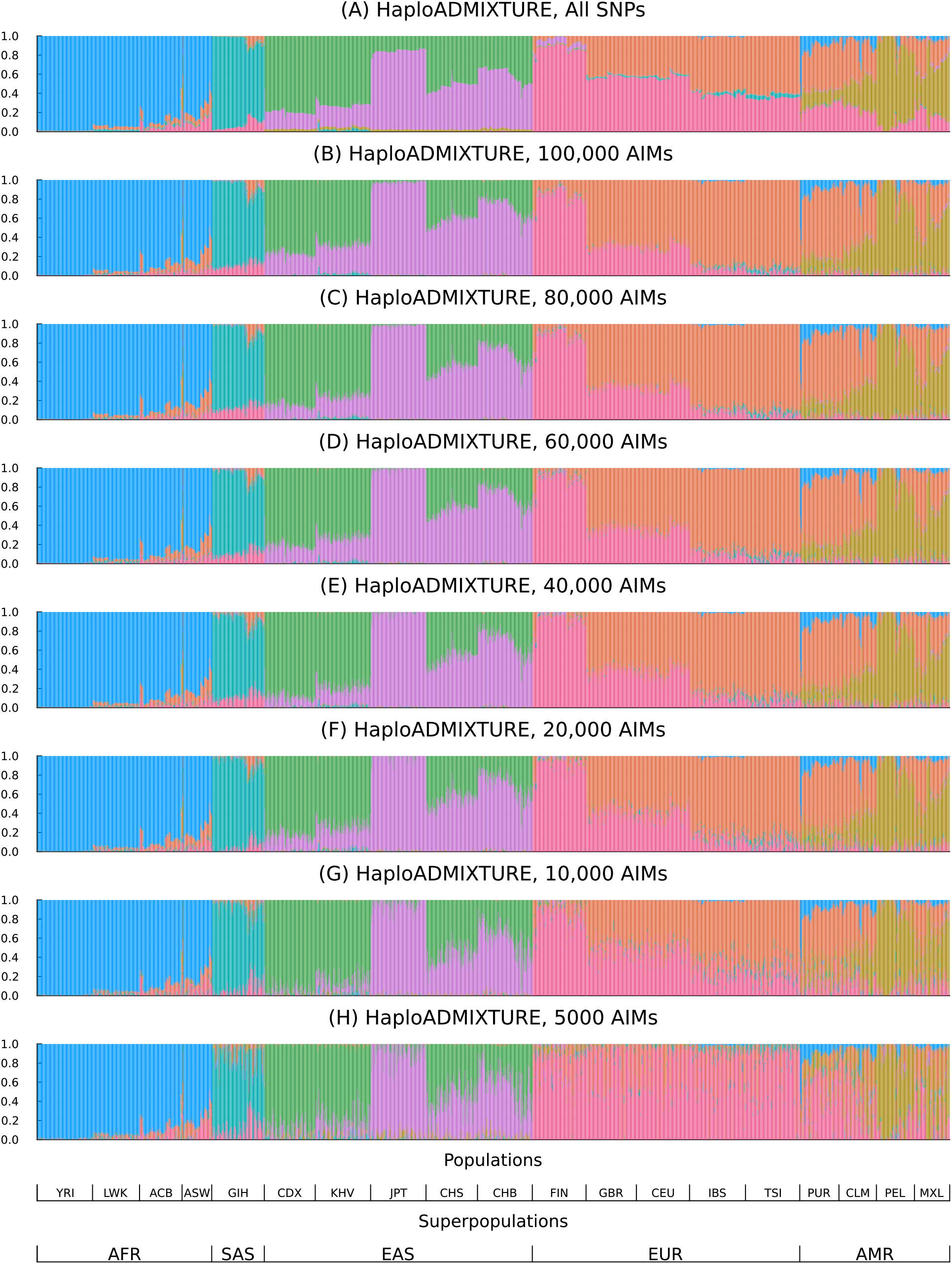
Ancestry estimation of TGP samples using different numbers of AIMs with HaploADMIXTURE. The results are presented in stacked bar plots, where the y-axis indicates the proportion of total ancestry. The x-axis shows all samples arranged by population labels.

**Figure 3.**
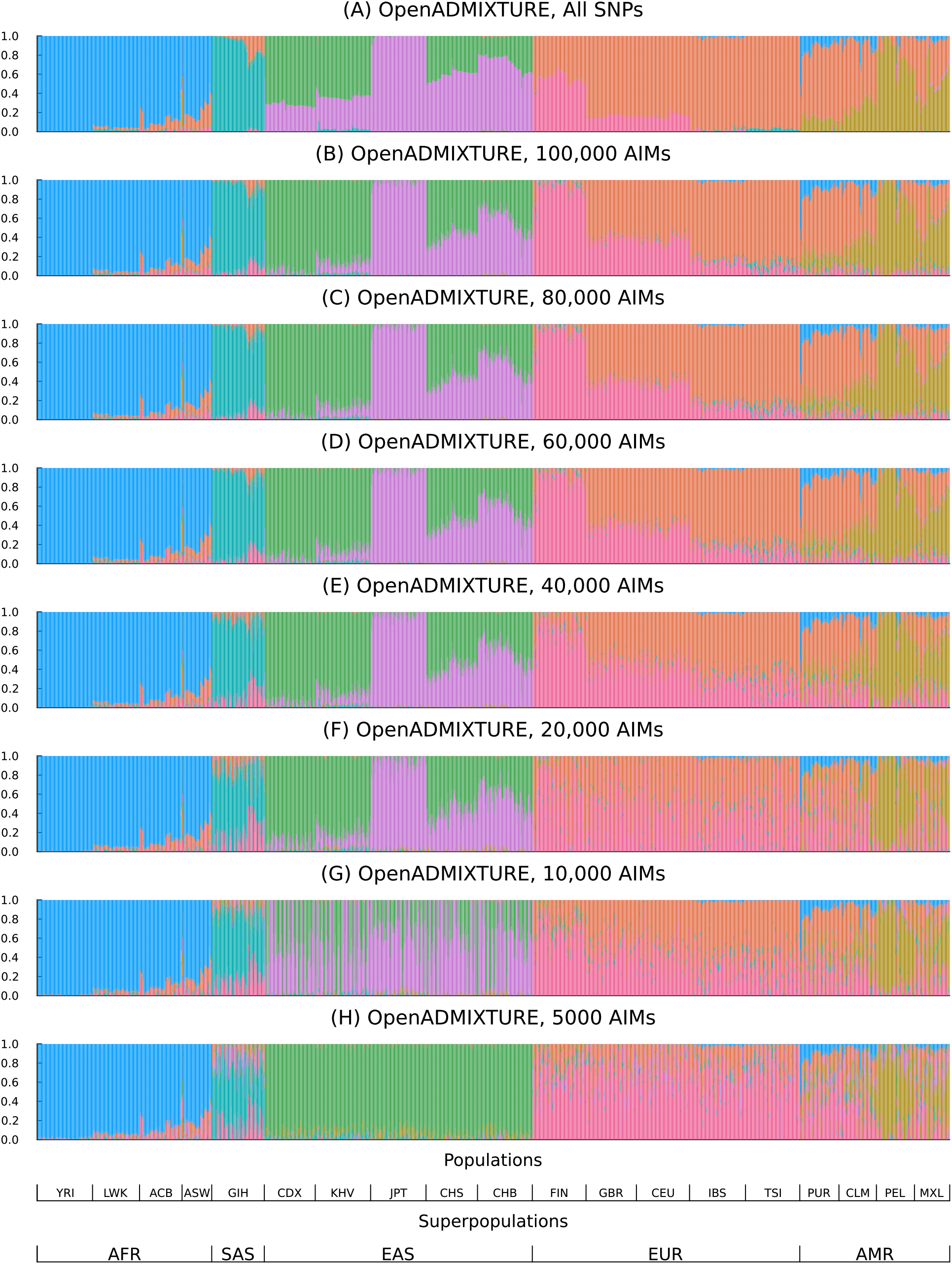
Ancestry estimation of TGP data samples using different numbers of AIMs with OpenADMIXTURE. The results are presented in stacked bar plots, where the y-axis indicates the proportion of total ancestry. The x-axis shows all samples arranged by population labels.

**Figure 4.**
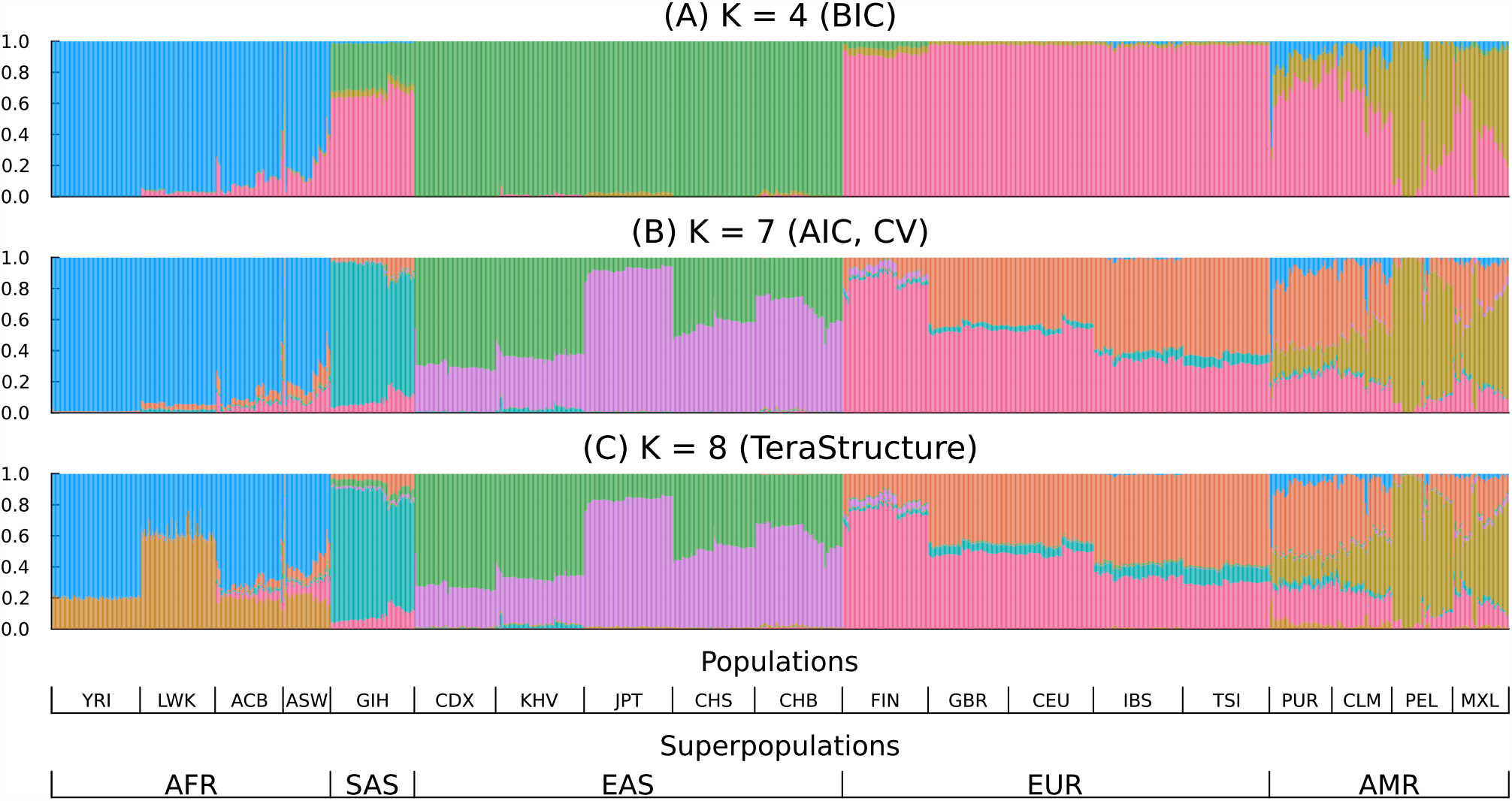
Structure inferred for TGP data samples using HaploADMIXTURE for different *K*. (a) *K* = 4 as selected by Bayesian information criterion, (b) *K* = 7 as selected by cross-validation and Akaike information criterion, (c) *K* = 8 as selected by the validation likelihood method in TeraStructure. The results are presented in stacked bar plots where the y-axis indicates the proportion of total ancestry. The x-axis shows all samples arranged by population labels.

#### 3.2.3 Loglikelihood and Entropy

Table S4 displays the likelihood of the fitted models. Since the binomial model of OpenADMIXTURE is a submodel of the model of HaploADMIXTURE, the maximum loglikelihood of the former is always less than the maximum loglikelihood of the latter based on the same SNP set. Table 2 shows the entropy of ***P***, the array of genotype/haplotype frequencies for each data set. The entropy decrease in HaploADMIXTURE compared to OpenADMIXTURE quantifies the additional information available in haplotypes. The entropy of ***P*** using HaploADMIXTURE for TGP, HGDP, and HO show 12.7%, 16.9%, and 18.9% reductions respectively, compared to OpenADMIXTURE. TeraStructure has higher entropy than OpenADMIXTURE, and SCOPE has entropy similar to OpenADMIXTURE on TGP and HO data sets. On HGDP, SCOPE has a similar entropy to HaploADMIXTURE. Note that the SCOPE model does not directly optimize the binomial loglikelihood model.

**Table 2.** Entropy per SNP per individual of P for TGP, HGDP, and HO.

| Software | TGP | HGDP | HO |
| --- | --- | --- | --- |
| HaploADMIXTURE | 0.303 | 0.344 | 0.360 |
| OpenADMIXTURE | 0.347 | 0.414 | 0.444 |
| SCOPE | 0.347 | 0.339 | 0.422 |
| TeraStructure | 0.393 | 0.432 | 0.474 |

#### 3.2.4 Evaluation of Estimated Admixture

Silhouette coefficients offer another way of quantifying performance. These are based on the ancestry labels implicit in the estimated ***Q*** matrix. The average silhouette coefficient is preferable to the training errors of linear classifiers and their cross-entropies^13,8^ because training error is discrete, and a single individual can unduly influence cross-entropy. We additionally matched clusters as discussed in Section 2.6.1 and computed root-mean-square error (RMSE) from the SKFR clusters derived from all SNPs.

Tables 4, S6, and S7 display mean silhouette coefficients for HaploADMIXTURE, OpenADMIXTURE, SCOPE, and TeraStructure. Since one of the continental populations is known to be admixed Americans, we also provide the result without them in Table S5. Tables S8–S10 show continent-by-continent mean silhouette coefficients, and Tables S11–S14 show region-by-region mean silhouette coefficients. HaploADMIXTURE generally performs well in grouping populations by both continent and region. OpenADMIXTURE performs equally well in grouping by continent but falls behind in grouping by region. For TGP and HGDP, TeraStructure is the best at distinguishing continental labels but falters in distinguishing regional labels, particularly in the TGP data where Middle-Eastern and European populations are lumped. For HGDP, SCOPE is the best at distinguishing the 32 regional labels but struggles compared to HaploADMIXTURE and OpenADMIXTURE in distinguishing African continental populations from each other. For the HO data set, HaploADMIXTURE and OpenADMIXTURE perform similarly in distinguishing the 163 regional labels, followed by SCOPE and TeraStructure.

When the analysis is based on AIMs, HaploADMIXTURE usually performs better than OpenADMIXTURE. In the single instance of 5000 AIMs for TGP, HaploADMIXTURE suffers in distinguishing regional subpopulations. In the case of HGDP under AIM selection, OpenADMIXTURE has trouble distinguishing between Middle-Eastern and European populations and adds a population to Africa. This anomaly is visible in Figure S4. HaploADMIXTURE with AIMs retains the power to distinguish the Middle-Eastern and European populations. For the HO data set, HaploADMIXTURE performance with AIMs better mimics its performance with all SNPs than OpenADMIXTURE does in the same comparison. Tables 3, S15, and S16 display RMSE from the baseline of all SNPs for the TGP, HGDP, and HO data sets, respectively. For the TGP data set, as we choose fewer AIMs, the mean silhouette tends to decrease, except for 10,000 and 5000 SNPs in OpenADMIXTURE. However, these exceptional cases yield poorer separation of populations than HaploADMIXTURE with all SNPs. This suggests that parsimony alone is an imperfect criterion for judging admixture estimation.

**Table 3.**
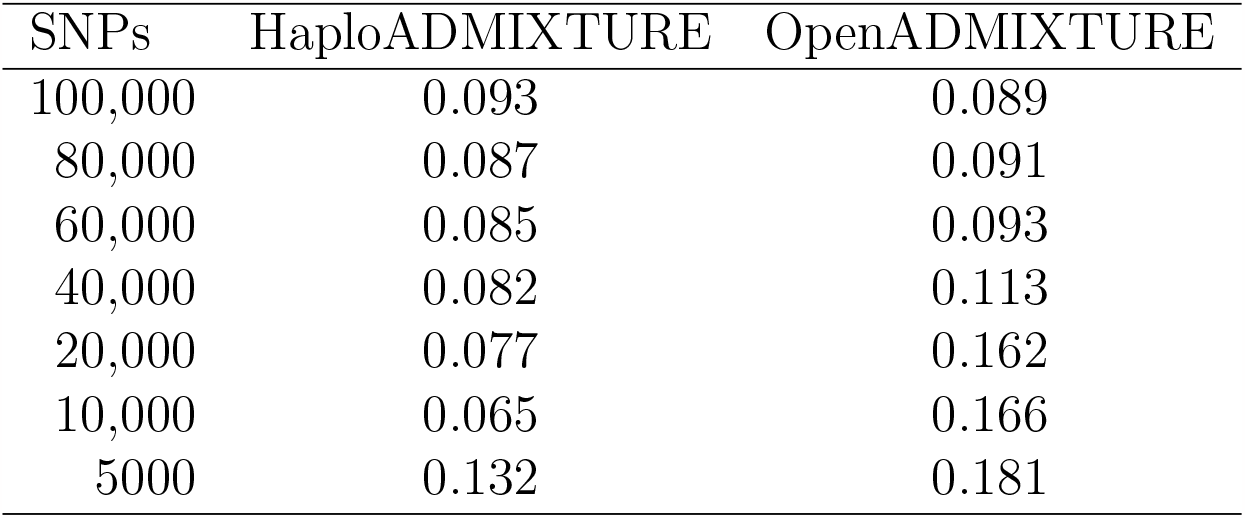
Root-mean-square error of sparse *K*-means (SKFR) from the baseline for HaploADMIXTURE and OpenADMIXTURE on the TGP data set. Root-meansquare error (RMSE) from baseline compares estimated admixture coefficients of SKFR to those estimated using all SNPs; the lower, the better.

| SNPs | HaploADMIXTURE | OpenADMIXTURE |
| --- | --- | --- |
| 100,000 | 0.093 | 0.089 |
| 80,000 | 0.087 | 0.091 |
| 60,000 | 0.085 | 0.093 |
| 40,000 | 0.082 | 0.113 |
| 20,000 | 0.077 | 0.162 |
| 10,000 | 0.065 | 0.166 |
| 5000 | 0.132 | 0.181 |

**Table 4.** Performance comparison of HaploADMIXTURE, OpenADMIXTURE, SCOPE, and TeraStructure on the TGP data set. Performance is measured by the mean silhouette coefficient of the population labels on the space of estimated admixture coefficients, **Q;** the higher, the better. The best value in the mean silhouette is in *italics*; these range over [−1 1].

| SNPs | HaploADMIXTURE | OpenADMIXTURE | SCOPE | TeraStructure |
| --- | --- | --- | --- | --- |
| Continental labels |  |  |  |  |
| 1,854,622 | 0.606 | 0.591 | 0.528 | *0.671* |
| 100,000 | 0.558 | 0.524 |  |  |
| 80,000 | 0.540 | 0.525 |  |  |
| 60,000 | 0.541 | 0.526 |  |  |
| 40,000 | 0.526 | 0.533 |  |  |
| 20,000 | 0.528 | 0.481 |  |  |
| 10,000 | 0.522 | 0.500 |  |  |
| 5000 | 0.521 | 0.626 |  |  |
| Regional labels |  |  |  |  |
| 1,854,622 | *0.423* | 0.413 | 0.418 | 0.335 |
| 100,000 | 0.360 | 0.347 |  |  |
| 80,000 | 0.353 | 0.329 |  |  |
| 60,000 | 0.335 | 0.296 |  |  |
| 40,000 | 0.317 | 0.225 |  |  |
| 20,000 | 0.277 | 0.147 |  |  |
| 10,000 | 0.206 | 0.035 |  |  |
| 5000 | 0.083 | 0.025 |  |  |

#### 3.3.5 Computational Efficiency

Given the computational improvements incorporated in HaploADMIXTURE, the analyses reported here finish in a reasonable amount of time. HaploADMIXTURE’s cost per iteration with *S* = 2 SNPs per haplotype block is less than eight times that of OpenADMIXTURE. Given that the number of frequency parameters quadruples, it takes four times longer to compute gradients and Hessians. While the time for solving quadratic programs is still negligible for ***Q***, quadratic programming for ***P*** takes longer, comparable to the time needed to compute gradients and Hessians on a GPU. Since 16-threaded ADMIXTURE was 16 times slower than GPU-accelerated OpenADMIXTURE^8^, HaploADMIXTURE’s per-iteration performance is still faster than that of ADMIXTURE. Balanced against these gains is the fact that the number of iterations until convergence increases. This reflects the greater complexity of the likelihood, the increased number of parameters, and the cost of parameter splitting by the MM principle.

Table S17 shows the average runtime using five random initial points for the TGP, HGDP, and HO data sets ignoring AIMs. Despite requiring more iterations to converge, HaploADMIXTURE takes less than 16 times longer than OpenADMIXTURE. Because runtime is proportional to the number of blocks *B* of SNPs employed, preprocessing with AIM selection to reduce *B* is recommended if speed is critical. For example, on the TGP data, it takes 2 minutes for sparse *K*-means to select 100,000 AIMs, and then another 12 minutes to run HaploADMIXTURE on the filtered dataset, for a total of just 14 minutes. Even so, running on AIMs yields admixture coefficients comparable to running on the full set of 1.8 million SNPs. The latter more onerous computations take 2 hours and 8 minutes. If one opts to preselect AIMs by sparse *K*-means, the time needed for SKFR in HaploADMIXTURE is not much different from that for OpenADMIXTURE. Indeed, the speed of the SKFR algorithm is minimally affected by the switch to haplotypes. SKFR and HaploADMIXTURE directly operate on PLINK BED-formatted data, so the total memory footprint of each is less than twice the size of the BED file.

## 4 Discussion

This paper introduces a technique for global ancestry estimation that converts linkage disequilibrium from a liability to an asset. Our program HaploADMIXTURE exploits multi-threading, GPU acceleration, and sparse *K*-means clustering to identify ancestry informative haplotypes. Although these advances also appear in OpenADMIXTURE, our earlier upgrade of the ADMIXTURE^6^ software, they require substantial modification to handle haplotypes. For instance, in the construction of AIMs, sparse *K*-means must now operate on haplotypes rather than SNPs. Likelihood calculation becomes more complicated because of increased phase ambiguity. Nevertheless, these technical hurdles can be overcome consistent with computational speed and memory demands on a par with or better than that of the original ADMIXTURE. Computation times scale linearly in the number of haplotype blocks. To keep computational costs in check, our haplotypes span just two SNPs. Even with this limitation, we see substantial gains in ancestry estimation precision. Extending haplotype blocks to include more than two SNPs is theoretically possible and would further increase information content, but extension quickly hits a combinatorial wall in computing the 2^*S*^ haplotype frequencies given *S* SNPs per block. The greater phase ambiguity encountered would complicate computer code and slow the convergence of recursive quadratic programming, the optimization engine in HaploADMIXTURE.

The admixture coefficients delivered by HaploADMIXTURE demonstrate a good separation of populations at the continental and regional levels in both real and simulated data sets. The other admixture programs tested often perform well on one level and poorly on the other. The admixture estimates from HaploADMIXTURE are more accurate than the competition as measured by mean square prediction error. In our experience, cross-validation and AIC produce reasonable estimates of the number of ancestral populations *K*. AIC is much faster than cross-validation. It will be interesting to see whether Bayesian or algebraic methods can be adapted to exploit haplotypes. The algebraic program SCOPE relies on alternating least squares. Adaptation would require passing from matrix to tensor decompositions.

Estimation of admixture proportions given known populations and known haplotype frequencies is possible with HaploADMIXTURE. One simply fixes ***P*** and updates only ***Q***. This simplification is invoked in the time-consuming process of cross-validation. For optimal general use, partial maximization would require curating the most informative pairs of SNPs in population panels. Partial maximization is a convex problem, with parameter separation across individuals, and thus easily applied to biobank-scale subjects.

In summary, HaploADMIXTURE is a thoughtful extension of OpenADMIXTURE, the open-source upgrade of ADMIXTURE. Modeling haplotypes adds vital information in ancestry analysis, yields more precise estimates of admixture coefficients, and distinguishes subpopulations better. HaploADMIXTURE builds on Julia’s high-performance computing environment and leverages potent OpenMendel tools. As HaploADMIXTURE is expanded and improved over time, we hope that it will ultimately receive the wide acceptance already enjoyed by ADMIXTURE.

## Supplemental Data

Supplemental data include nine figures and 11 tables.

## Acknowledgments

This research was partially funded by grants from the National Institute of General Medical Sciences (R35GM141798, EMS, HZ, and KL), the National Human Genome Research Institute (R01HG006139, EMS, HZ, and KL), and the National Science Foundation (DMS-2054253 and IIS-2205441, HZ).

## Web Resources

OpenADMIXTURE, https://github.com/OpenMendel/OpenADMIXTURE.jl.

SKFR, https://github.com/kose-y/SKFR.jl.

SnpArrays, https://github.com/OpenMendel/SnpArrays.jl.

SCOPE, https://github.com/sriramlab/SCOPE.

## Data and Code Availability

The HaploADMIXTURE package can be found at https://github.com/OpenMendel/HaploADMIXTURE.jl. The code for the experiments and instructions to download publicly available data can be found at https://github.com/kose-y/HaploADMIXTURE-resources.

