## Supplemental Material for "Estimation of Genetic Admixture Proportions via Haplotypes"

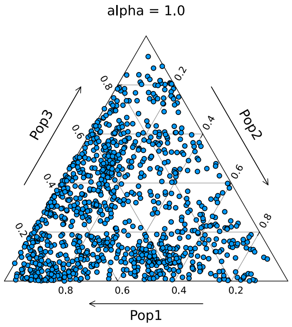


Figure S1: **Visualization of dispersion of $\boldsymbol{Q}$ with $K = 3$ and varying values of $\alpha$.**

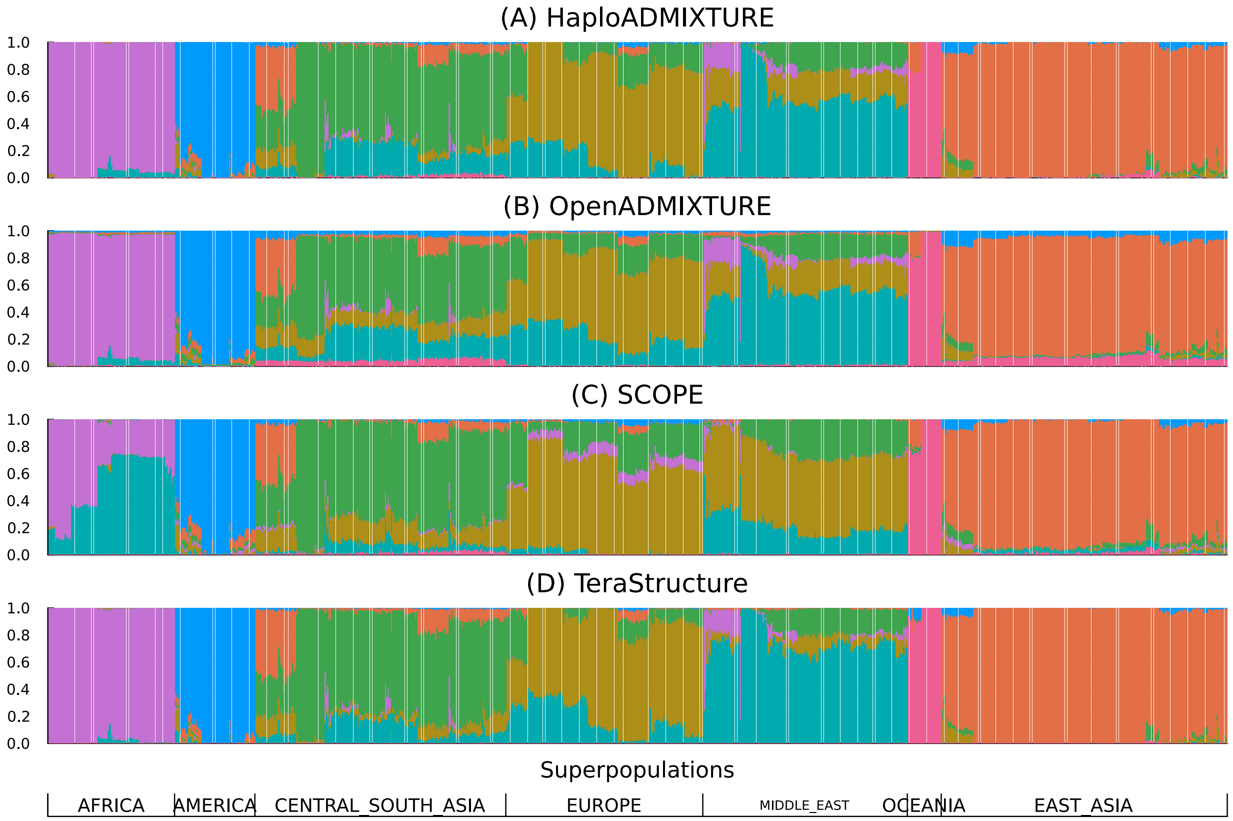


Figure S2: **Ancestry estimation of HGDP data samples**. (a) Using HaploADMIXTURE with all SNPs, (b) OpenADMIXTURE with all SNPs, (c) SCOPE, and (d) TeraStructure. The results are presented in stacked bar plots, where the y-axis indicates the proportion of total ancestry. The x-axis shows all samples arranged by population labels.

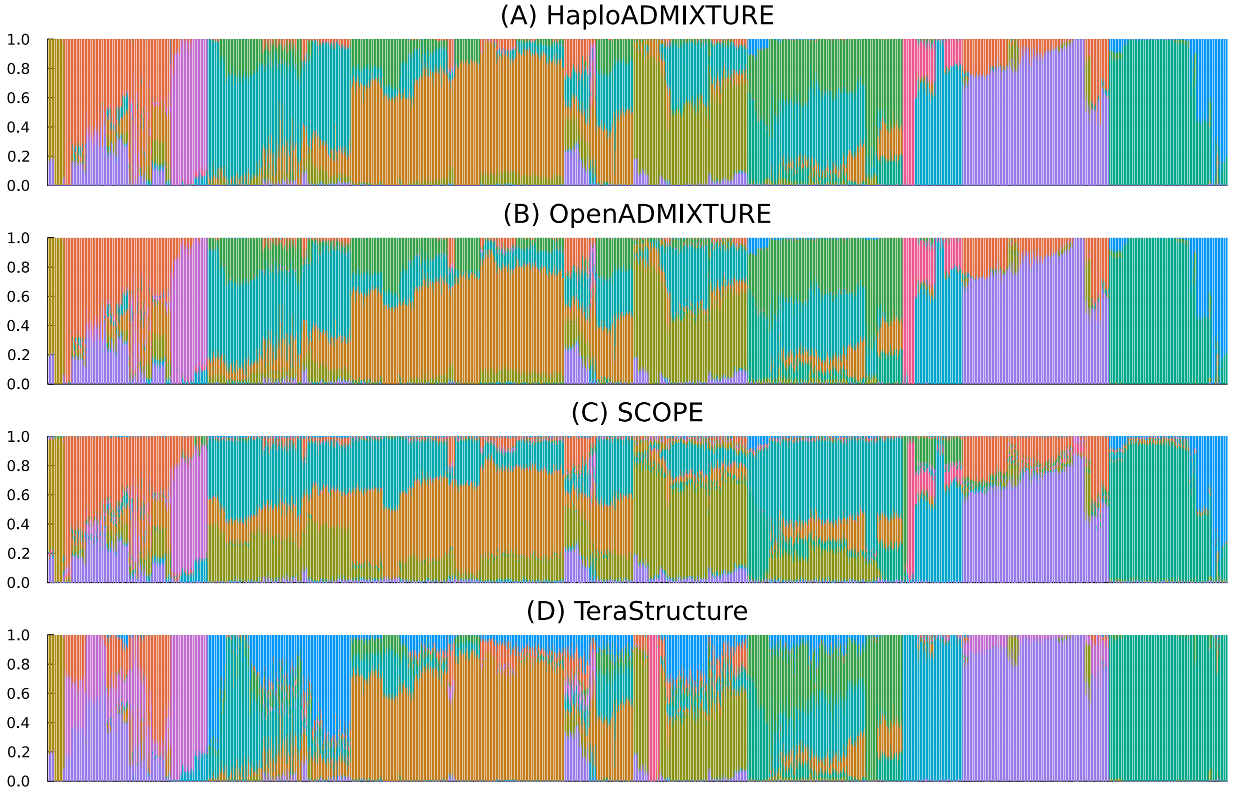


Figure S3: **Ancestry estimation of HO data samples**. (a) Using HaploADMIXTURE with all SNPs, (b) OpenADMIXTURE with all SNPs, (c) SCOPE, and (d) TeraStructure. The results are presented in stacked bar plots, where the y-axis indicates the proportion of total ancestry. The x-axis shows all the samples arranged by population labels.

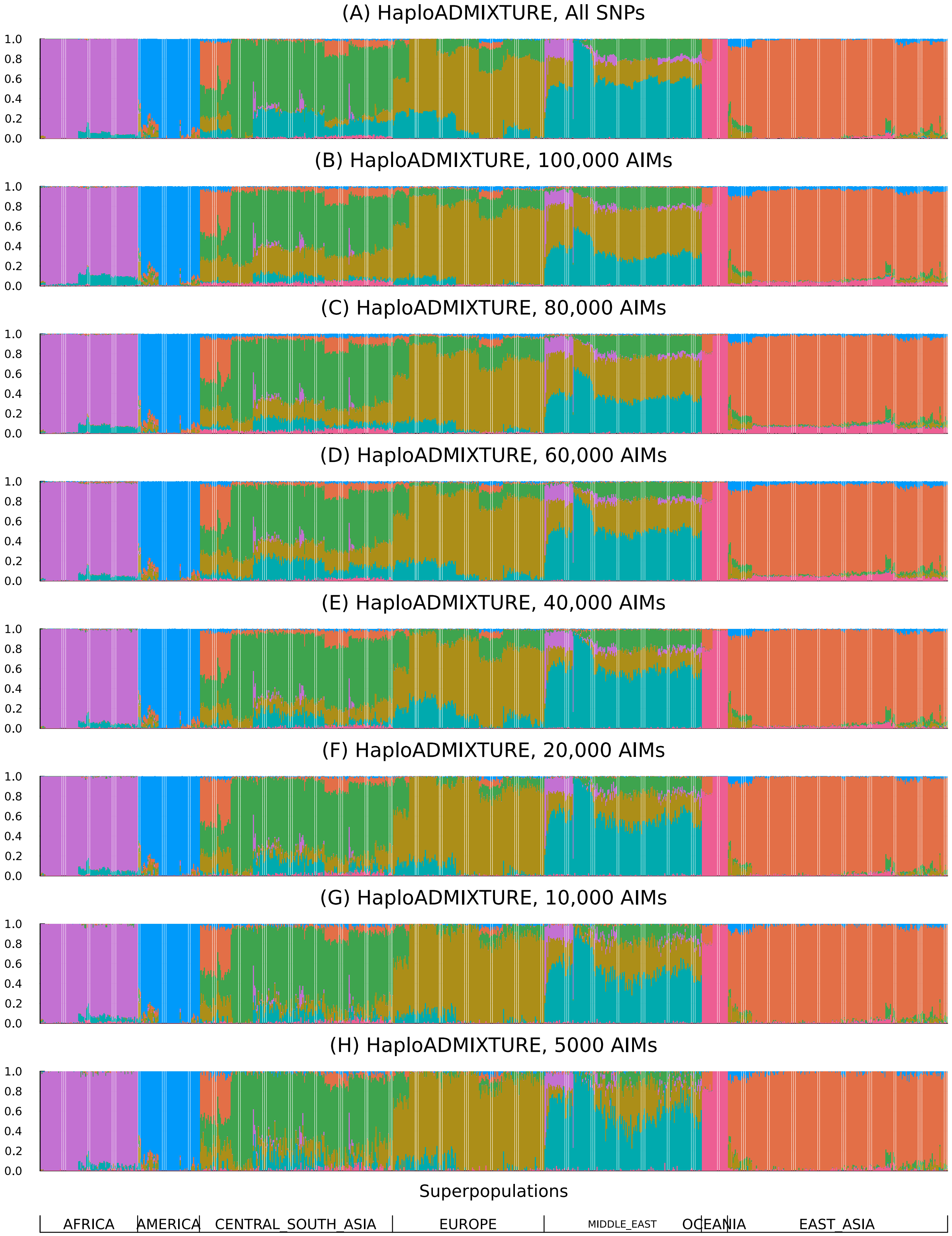


Figure S4: **Ancestry estimation of HGDP data samples using different numbers of AIMs on HaploADMIXTURE**. The results are presented in stacked bar plots, where the y-axis indicates the proportion of total ancestry. The x-axis shows all samples arranged by population labels.

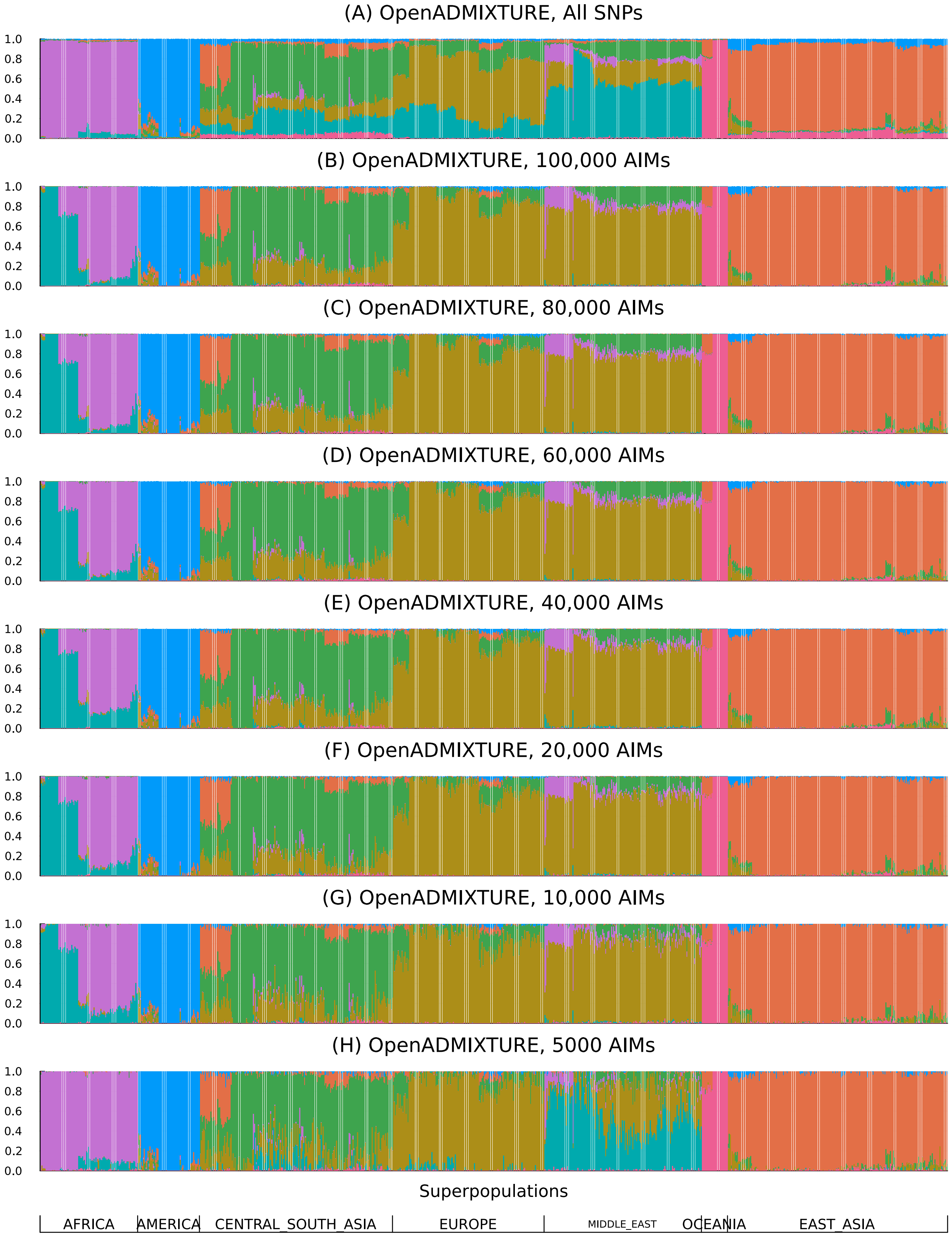


Figure S5: **Ancestry estimation of HGDP data samples using different numbers of AIMs on OpenADMIXTURE**. The results are presented in stacked bar plots, where the y-axis indicates the proportion of total ancestry. The x-axis shows all samples arranged by population labels.

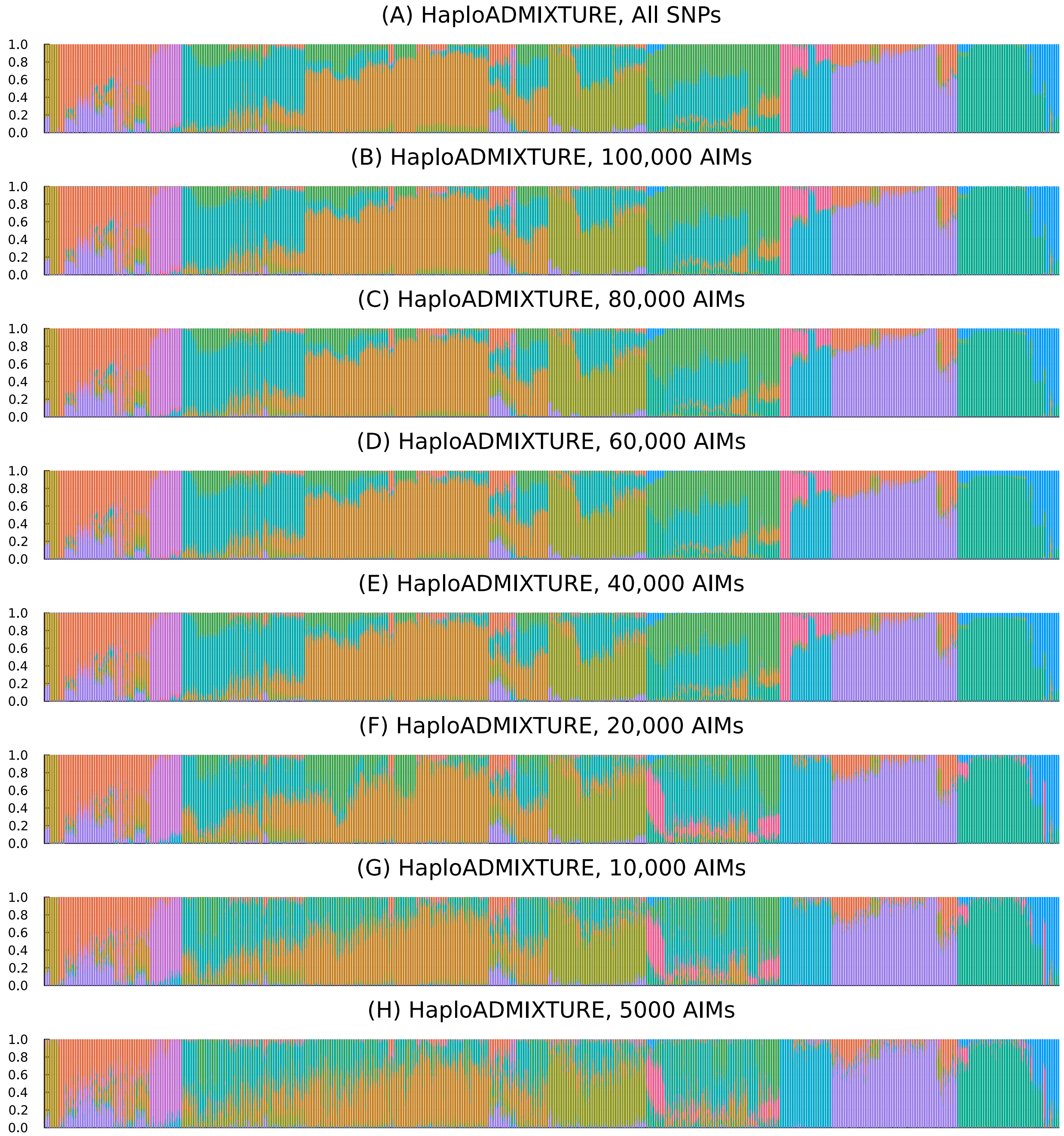


Figure S6: **Ancestry estimation of HO data samples using different numbers of AIMs on HaploADMIXTURE**. The results are presented in stacked bar plots, where the y-axis indicates the proportion of total ancestry. The x-axis shows all samples arranged by population labels.

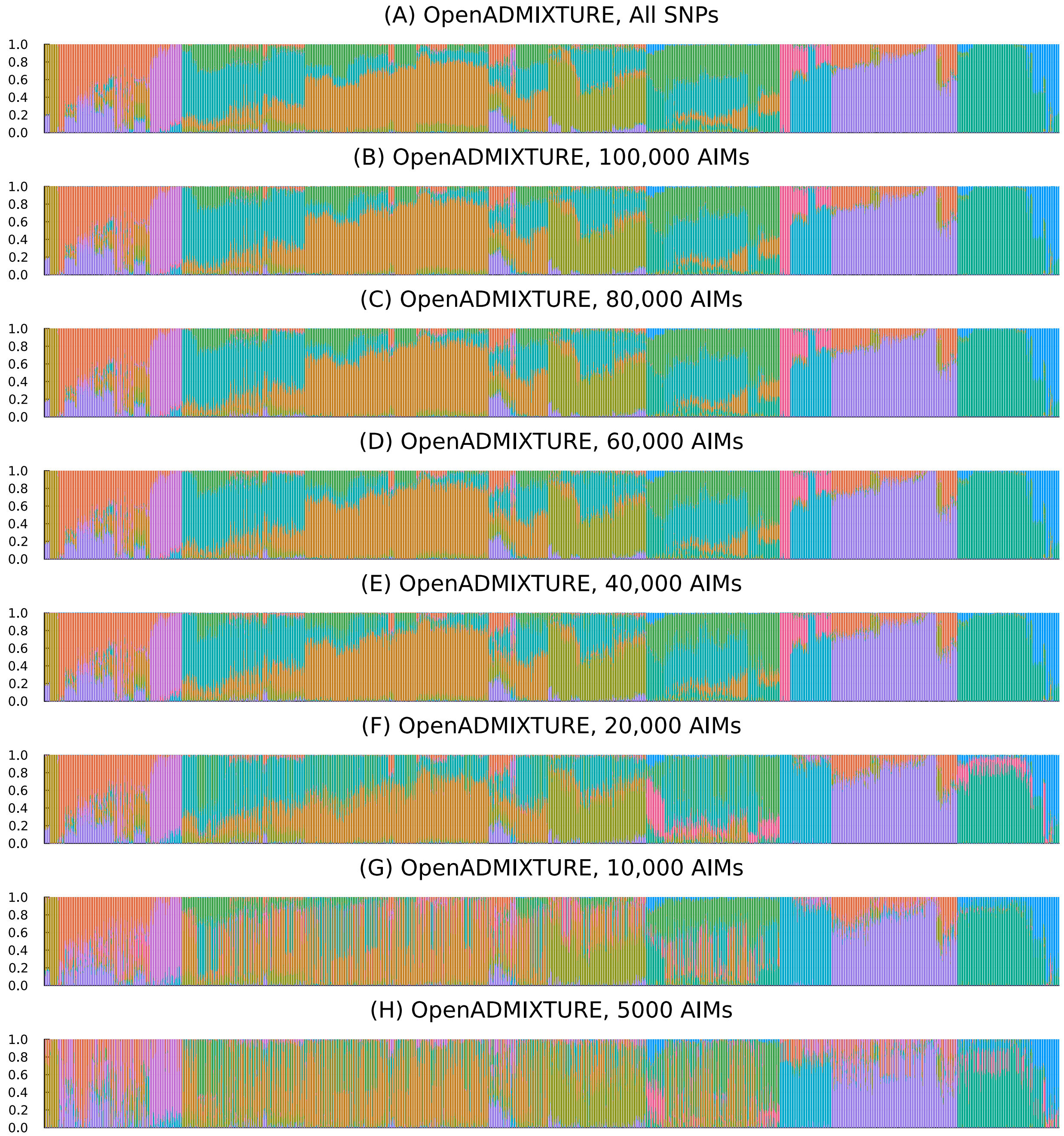


Figure S7: **Ancestry estimation of HO data samples using different numbers of AIMs on OpenADMIXTURE**. The results are presented in stacked bar plots where the y-axis indicates the proportion of total ancestry. The x-axis shows all samples arranged by population labels.

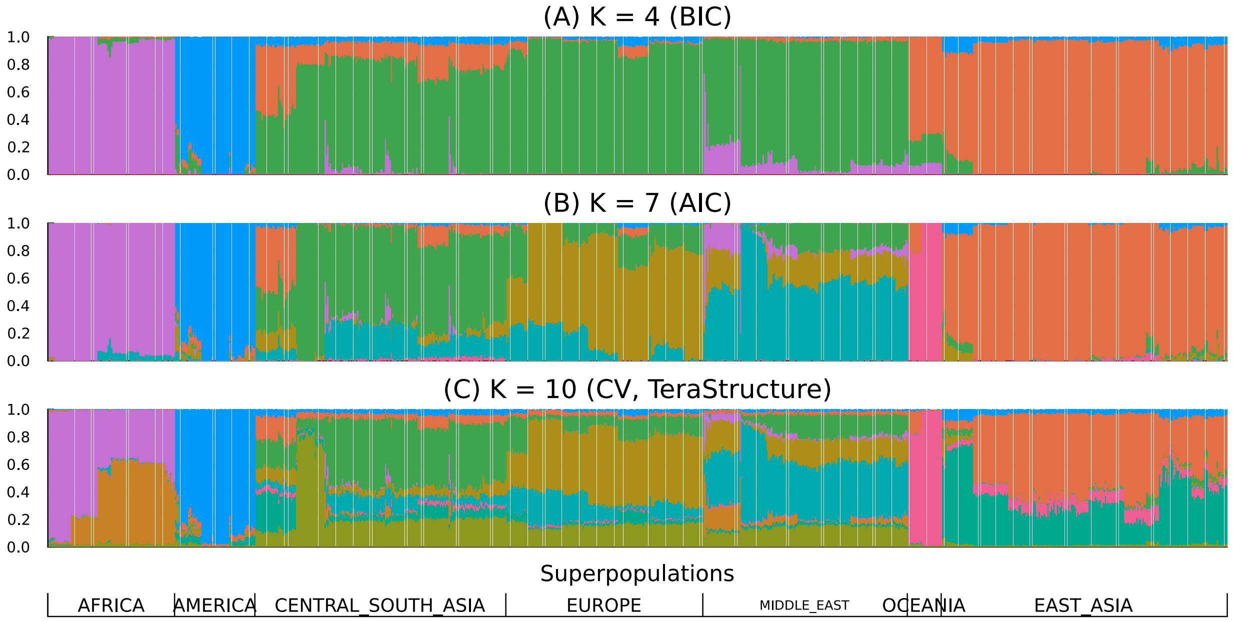


Figure S8: **Structure inferred for HGDP data samples using HaploADMIXTURE for different** $K$. (a) $K = 7$ chosen by AIC, (b) $K = 10$ chosen by the AIC, and the validation likelihood method in TeraStructure. The results are presented in stacked bar plots, where the y-axis indicates the proportion of total ancestry. The x-axis shows all samples arranged by population labels.

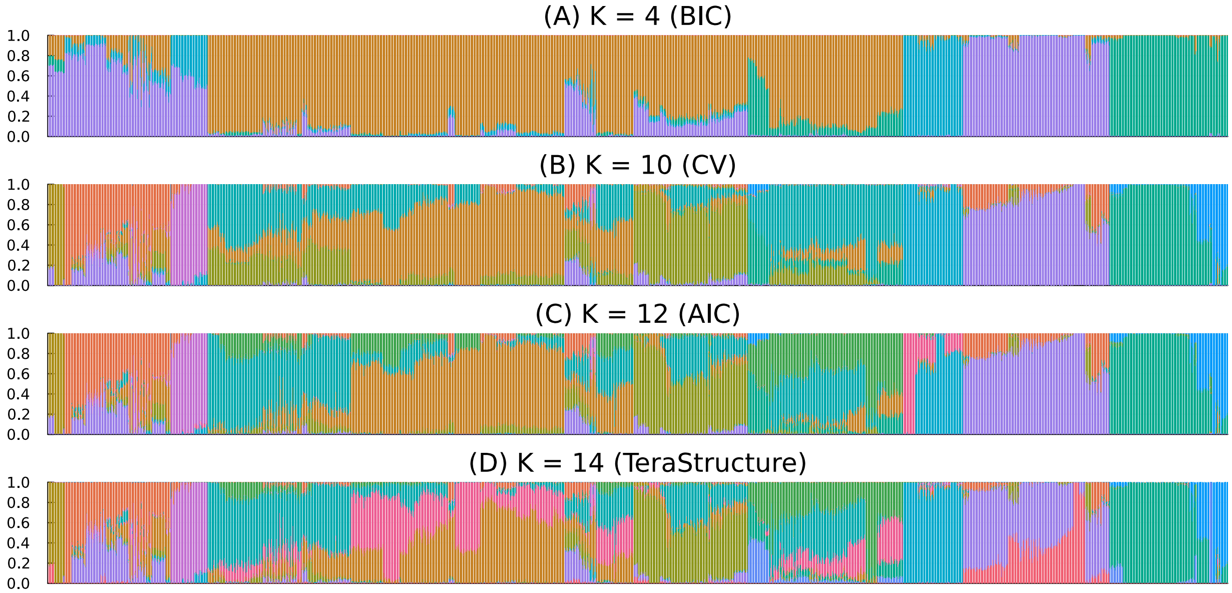


Figure S9: **Structure inferred for HO data samples using HaploADMIXTURE for different** $K$. (a) $K = 10$ chosen by CV, (b) $K = 12$ chosen by the AIC, and (c) $K = 14$ chosen by the validation likelihood method in TeraStructure. The results are presented in stacked bar plots, where the y-axis indicates the proportion of total ancestry. The x-axis shows all samples arranged by population labels.

Table S1: **Weights $w^{(n)}_{ibk\boldsymbol{h}}$ corresponding to genotype $\boldsymbol{x}_{il}$ for $S = 2$.** An asterisk $(*)$ denotes a missing genotype, and $v_{\boldsymbol{h}}$ denotes $\sum_k q^{(n)}_{ki} p^{(n)}_{kb\boldsymbol{h}}$, where $q^{(n)}_{ki}$ and $p^{(n)}_{kb\boldsymbol{h}}$ are the current iterates for $q_{ki}$ and $p_{kb\boldsymbol{h}}$, respectively.

| Genotype $\boldsymbol{x}_{il}$ | $w^{(n)}_{ibk(00)}$ | $w^{(n)}_{ibk(01)}$ | $w^{(n)}_{ibk(10)}$ | $a^{(n)}_{ilk(11)}$ |
|---|---|---|---|---|
| $(**)$ | 0 | 0 | 0 | 0 |
| $(*0)$ | $\frac{2v_{(00)}}{v_{(00)}+v_{(10)}}$ | 0 | $\frac{2v_{(10)}}{v_{(00)}+v_{(10)}}$ | 0 |
| $(*1)$ | $\frac{v_{(00)}}{v_{(00)}+v_{(10)}}$ | $\frac{v_{(01)}}{v_{(01)}+v_{(11)}}$ | $\frac{v_{(10)}}{v_{(00)}+v_{(10)}}$ | $\frac{v_{(11)}}{v_{(01)}+v_{(11)}}$ |
| $(*2)$ | 0 | $\frac{2v_{(01)}}{v_{(01)}+v_{(11)}}$ | 0 | $\frac{2v_{(11)}}{v_{(01)}+v_{(11)}}$ |
| $(0*)$ | $\frac{2v_{(00)}}{v_{(00)}+v_{(01)}}$ | $\frac{2v_{(01)}}{v_{(00)}+v_{(01)}}$ | 0 | 0 |
| $(1*)$ | $\frac{v_{(00)}}{v_{(00)}+v_{(01)}}$ | $\frac{v_{(01)}}{v_{(00)}+v_{(01)}}$ | $\frac{v_{(10)}}{v_{(10)}+v_{(11)}}$ | $\frac{v_{(11)}}{v_{(10)}+v_{(11)}}$ |
| $(2*)$ | 0 | 0 | $\frac{2v_{(10)}}{v_{(10)}+v_{(11)}}$ | $\frac{2v_{(11)}}{v_{(10)}+v_{(11)}}$ |
| (00) | 2 | 0 | 0 | 0 |
| (01) | 1 | 1 | 0 | 0 |
| (02) | 0 | 2 | 0 | 0 |
| (10) | 1 | 0 | 1 | 0 |
| (11) | $\frac{v_{(00)}v_{(11)}}{v_{(00)}v_{(11)}+v_{(01)}v_{(10)}}$ | $\frac{v_{(01)}v_{(10)}}{v_{(00)}v_{(11)}+v_{(01)}v_{(10)}}$ | $\frac{v_{(01)}v_{(10)}}{v_{(00)}v_{(11)}+v_{(01)}v_{(10)}}$ | $\frac{v_{(00)}v_{(11)}}{v_{(00)}v_{(11)}+v_{(01)}v_{(10)}}$ |
| (12) | 0 | 1 | 0 | 1 |
| (20) | 0 | 0 | 2 | 0 |
| (21) | 0 | 0 | 1 | 1 |
| (22) | 0 | 0 | 0 | 2 |

Table S2: **Root-mean-square errors of the estimated admixture proportions on the simulated data sets.** Root-mean-square error checks the accuracy of the estimated admixture coefficients; the lower, the better. Five populations were used for the simulation, with 1000 individuals and 1,000,000 SNPs for various values of $\alpha$. Each value is averaged over five simulation runs. The best value for each $\alpha$ is in *italics*.

| AIMs | HaploADMIXTURE | OpenADMIXTURE | SCOPE | TeraStructure |
|---|---|---|---|---|
| 1,000,000 SNPs, $\alpha = 0.1$ | | | | |
| 1,000,000 | 0.0060 | *0.0026* | 0.0060 | 0.0103 |
| 200,000 | 0.0042 | 0.0037 | | |
| 150,000 | 0.0042 | 0.0039 | | |
| 100,000 | 0.0042 | 0.0042 | | |
| 50,000 | 0.0045 | 0.0050 | | |
| 1,000,000 SNPs, $\alpha = 0.05$ | | | | |
| 1,000,000 | 0.0029 | 0.0027 | 0.0083 | 0.0069 |
| 200,000 | *0.0024* | 0.0037 | | |
| 150,000 | 0.0025 | 0.0040 | | |
| 100,000 | 0.0027 | 0.0044 | | |
| 50,000 | 0.0033 | 0.0055 | | |
| 1,000,000 SNPs, $\alpha = 0.02$ | | | | |
| 1,000,000 | 0.0016 | 0.0030 | 0.0101 | 0.0063 |
| 200,000 | *0.0015* | 0.0043 | | |
| 150,000 | 0.0017 | 0.0045 | | |
| 100,000 | 0.0020 | 0.0050 | | |
| 50,000 | 0.0028 | 0.0059 | | |

Table S3: **Root-mean-square errors of estimated admixture proportion on the simulated data sets.** Root-mean-square error checks the accuracy of the estimated admixture coefficients; the lower, the better. Five populations were used for the simulation, with 1000 individuals and 100,000 SNPs with various values of $\alpha$. Each value is averaged over five simulation runs. The best value for each $\alpha$ is in italics.

| AIMs | HaploADMIXTURE | OpenADMIXTURE | SCOPE | TeraStructure |
|---|---|---|---|---|
| 100,000 SNPs, $\alpha = 0.1$ | | | | |
| 100,000 | *0.0077* | 0.0083 | 0.0192 | 0.0253 |
| 20,000 | 0.0083 | 0.0110 | | |
| 15,000 | 0.0090 | 0.0119 | | |
| 10,000 | 0.0101 | 0.0132 | | |
| 5000 | 0.0127 | 0.0160 | | |
| 100,000 SNPs, $\alpha = 0.05$ | | | | |
| 100,000 | *0.0050* | 0.0100 | 0.0230 | 0.0270 |
| 20,000 | 0.0073 | 0.0126 | | |
| 15,000 | 0.0080 | 0.0133 | | |
| 10,000 | 0.0092 | 0.0143 | | |
| 5000 | 0.0117 | 0.0164 | | |
| 100,000 SNPs, $\alpha = 0.02$ | | | | |
| 100,000 | *0.0039* | 0.0111 | 0.0254 | 0.0288 |
| 20,000 | 0.0070 | 0.0134 | | |
| 15,000 | 0.0077 | 0.0140 | | |
| 10,000 | 0.0088 | 0.0150 | | |
| 5000 | 0.0111 | 0.0167 | | |

Table S4: **Loglikelihood of the fitted models for TGP, HGDP, and HO.** Since the HaploADMIXTURE model includes the OpenADMIXTURE model, HaploADMIXTURE shows a higher loglikelihood.

| Software | TGP | HGDP | HO |
|---|---|---|---|
| HaploADMIXTURE | -2.036E+09 | -4.587E+08 | -5.784E+08 |
| OpenADMIXTURE | -2.320E+09 | -5.349E+08 | -7.201E+08 |
| SCOPE | -2.322E+09 | -5.360E+08 | -7.208E+08 |
| TeraStructure | -2.326E+09 | -5.357E+08 | -7.225E+08 |

Table S5: **Performance comparison of HaploADMIXTURE, OpenADMIXTURE, SCOPE, and TeraStructure on the TGP data set without Admixed American population.** Performance is measured by the mean silhouette coefficient of the population labels on the space of estimated admixture coefficients, $\boldsymbol{Q}$; the higher, the better. The best value in the mean silhouette is in *italics*; these range over $[-1, 1]$.

| SNPs | HaploADMIXTURE | OpenADMIXTURE | SCOPE | TeraStructure |
|---|---|---|---|---|
| | Continental labels | | | |
| 1,854,622 | 0.740 | 0.749 | 0.664 | *0.805* |
| 100,000 | 0.694 | 0.661 | | |
| 80,000 | 0.671 | 0.663 | | |
| 60,000 | 0.672 | 0.664 | | |
| 40,000 | 0.652 | 0.668 | | |
| 20,000 | 0.666 | 0.598 | | |
| 10,000 | 0.657 | 0.635 | | |
| 5000 | 0.658 | 0.764 | | |
| | Regional labels | | | |
| 1,854,622 | *0.474* | 0.464 | 0.468 | 0.371 |
| 100,000 | 0.404 | 0.391 | | |
| 80,000 | 0.396 | 0.370 | | |
| 60,000 | 0.375 | 0.333 | | |
| 40,000 | 0.354 | 0.255 | | |
| 20,000 | 0.310 | 0.164 | | |
| 10,000 | 0.231 | 0.036 | | |
| 5000 | 0.100 | 0.028 | | |

Table S6: **Performance comparison of HaploADMIXTURE, OpenADMIXTURE, SCOPE, and TeraStructure on the HGDP data set.** Performance is measured by the mean silhouette coefficient of the population labels on the space of estimated admixture coefficients, $\boldsymbol{Q}$; the higher, the better. The best value in the mean silhouette is in *italics*; these range over $[-1, 1]$.

| SNPs | HaploADMIXTURE | OpenADMIXTURE | SCOPE | TeraStructure |
|---|---|---|---|---|
| | Continental labels | | | |
| 642,950 | 0.738 | 0.742 | 0.626 | *0.783* |
| 100,000 | 0.741 | 0.556 | | |
| 80,000 | 0.741 | 0.556 | | |
| 60,000 | 0.756 | 0.553 | | |
| 40,000 | 0.761 | 0.553 | | |
| 20,000 | 0.765 | 0.535 | | |
| 10,000 | 0.755 | 0.520 | | |
| 5000 | 0.744 | 0.697 | | |
| | Regional labels | | | |
| 642,950 | 0.258 | 0.224 | *0.298* | 0.131 |
| 100,000 | 0.223 | 0.174 | | |
| 80,000 | 0.200 | 0.167 | | |
| 60,000 | 0.171 | 0.144 | | |
| 40,000 | 0.141 | 0.104 | | |
| 20,000 | 0.091 | 0.050 | | |
| 10,000 | 0.049 | -0.003 | | |
| 5000 | -0.016 | -0.057 | | |

Table S7: **Performance comparison of HaploADMIXTURE, OpenADMIXTURE, SCOPE, and TeraStructure on the HO data set.** Performance is measured by the mean silhouette coefficient of the population labels on the space of estimated admixture coefficients, $\boldsymbol{Q}$; the higher, the better. The best value in the mean silhouette is in *italics*; these range over $[-1, 1]$.

| SNPs | HaploADMIXTURE | OpenADMIXTURE | SCOPE | TeraStructure |
|---|---|---|---|---|
| | Population labels | | | |
| 385,088 | *0.169* | *0.170* | 0.140 | 0.045 |
| 100,000 | 0.100 | 0.076 | | |
| 80,000 | 0.081 | 0.055 | | |
| 60,000 | 0.067 | 0.029 | | |
| 40,000 | 0.023 | -0.013 | | |
| 20,000 | -0.057 | -0.165 | | |
| 10,000 | -0.156 | -0.243 | | |
| 5000 | -0.224 | -0.352 | | |

Table S8: **Continent-by-continent mean silhouette for the TGP data set.** These represent averages over each population label on the space of estimated admixture coefficients, $\boldsymbol{Q}$; the higher, the better. Population labels in "Closest" indicate the continent closest to the label in the "Continent" column.

| | HaploADMIXTURE | | OpenADMIXTURE | | SCOPE | | TeraStructure | |
|---|---|---|---|---|---|---|---|---|
| Continent | Silhouette | Closest | Silhouette | Closest | Silhouette | Closest | Silhouette | Closest |
| AFR | 0.871 | AMR | 0.870 | AMR | 0.862 | AMR | 0.871 | AMR |
| AMR | 0.273 | EUR | 0.180 | EUR | 0.274 | EUR | 0.104 | EUR |
| EAS | 0.616 | AMR | 0.618 | AMR | 0.498 | AMR | 0.543 | AMR |
| EUR | 0.558 | AMR | 0.561 | AMR | 0.417 | AMR | 0.954 | AMR |
| SAS | 0.876 | AMR | 0.852 | AMR | 0.857 | AMR | 0.835 | AMR |

Table S9: **Continent-by-continent mean silhouette for the TGP data set without admixed American population.** These represent averages over each population label on the space of estimated admixture coefficients, $\boldsymbol{Q}$; the higher, the better.

| | HaploADMIXTURE | | OpenADMIXTURE | | SCOPE | | TeraStructure | |
|---|---|---|---|---|---|---|---|---|
| Continent | Silhouette | Closest | Silhouette | Closest | Silhouette | Closest | Silhouette | Closest |
| AFR | 0.882 | EUR | 0.881 | EUR | 0.873 | EUR | 0.884 | EUR |
| EAS | 0.639 | EUR | 0.645 | EUR | 0.524 | EUR | 0.576 | SAS |
| EUR | 0.722 | EAS | 0.747 | EAS | 0.629 | SAS | 0.975 | SAS |
| SAS | 0.877 | EUR | 0.855 | EUR | 0.860 | EUR | 0.842 | EUR |

Table S10: **Continent-by-continent mean silhouette for the HGDP data set.** These represent averages over each population label on the space of estimated admixture coefficients, $\boldsymbol{Q}$, the higher, the better. AFR: Africa, AMR: America, CSA: Central South Asia, EAS: East Asia, EUR: Europe, ME: Middle East, OCE: Oceania.

| | HaploADMIXTURE | | OpenADMIXTURE | | SCOPE | | TeraStructure | |
|---|---|---|---|---|---|---|---|---|
| Continent | Silhouette | Closest | Silhouette | Closest | Silhouette | Closest | Silhouette | Closest |
| AFR | 0.949 | ME | 0.954 | ME | 0.603 | ME | 0.973 | ME |
| AMR | 0.881 | CSA | 0.886 | ME | 0.858 | ME | 0.904 | CSA |
| CSA | 0.552 | ME | 0.556 | ME | 0.580 | ME | 0.600 | ME |
| EAS | 0.890 | CSA | 0.899 | CSA | 0.892 | CSA | 0.924 | CSA |
| EUR | 0.634 | ME | 0.629 | ME | 0.368 | ME | 0.650 | ME |
| ME | 0.644 | EUR | 0.669 | EUR | 0.444 | EUR | 0.751 | CSA |
| OCE | 0.873 | CSA | 0.859 | CSA | 0.851 | ME | 0.936 | CSA |

Table S11: **Region-by-region mean silhouette for the TGP data set.** These represent averages over each population label on the space of estimated admixture coefficients, $\boldsymbol{Q}$, the higher, the better.

| | HaploADMIXTURE | | OpenADMIXTURE | | SCOPE | | TeraStructure | | |
|---|---|---|---|---|---|---|---|---|---|
| Population | Silhouette | Closest | Silhouette | Closest | Silhouette | Closest | Silhouette | Closest | Continent |
| ACB | -0.213 | ASW | -0.188 | ASW | -0.030 | ASW | -0.216 | ASW | AFR |
| ASW | -0.112 | ACB | -0.112 | ACB | -0.091 | ACB | -0.109 | ACB | AFR |
| LWK | 0.703 | YRI | 0.636 | YRI | 0.712 | YRI | 0.229 | YRI | AFR |
| YRI | 0.927 | LWK | 0.989 | LWK | 0.783 | LWK | 1.000 | LWK | AFR |
| CLM | 0.044 | IBS | 0.008 | IBS | 0.030 | IBS | 0.009 | FIN | AMR |
| MXL | -0.145 | IBS | -0.138 | IBS | -0.105 | IBS | -0.169 | FIN | AMR |
| PEL | 0.467 | MXL | 0.435 | MXL | 0.474 | MXL | 0.518 | MXL | AMR |
| PUR | 0.274 | CLM | 0.280 | CLM | 0.239 | IBS | 0.251 | FIN | AMR |
| CDX | 0.592 | KHV | 0.600 | KHV | 0.534 | KHV | 0.541 | KHV | EAS |
| CHB | 0.205 | CHS | 0.208 | CHS | 0.211 | CHS | 0.147 | CHS | EAS |
| CHS | 0.585 | CHB | 0.580 | CHB | 0.576 | CHB | 0.574 | CHB | EAS |
| JPT | 0.913 | CHB | 0.963 | CHB | 0.893 | CHB | 0.983 | CHB | EAS |
| KHV | 0.543 | CDX | 0.555 | CDX | 0.529 | CDX | 0.563 | CDX | EAS |
| CEU | -0.118 | GBR | -0.151 | GBR | -0.080 | GBR | -0.306 | GBR | EUR |
| FIN | 0.826 | CEU | 0.805 | CEU | 0.798 | CEU | 0.631 | TSI | EUR |
| GBR | 0.127 | CEU | 0.165 | CEU | 0.081 | CEU | 0.375 | CEU | EUR |
| IBS | 0.282 | TSI | 0.071 | TSI | 0.381 | TSI | -0.383 | TSI | EUR |
| TSI | 0.560 | IBS | 0.596 | IBS | 0.514 | IBS | 0.346 | CEU | EUR |
| GIH | 0.867 | PUR | 0.845 | CLM | 0.850 | CLM | 0.824 | CLM | SAS |

Table S12: **Region-by-region mean silhouette for the TGP data set without admixed American population.** These represent averages over each population label on the space of estimated admixture coefficients, $\boldsymbol{Q}$, the higher, the better.

| | HaploADMIXTURE | | OpenADMIXTURE | | SCOPE | | TeraStructure | | |
|---|---|---|---|---|---|---|---|---|---|
| Population | Silhouette | Closest | Silhouette | Closest | Silhouette | Closest | Silhouette | Closest | Continent |
| ACB | -0.213 | ASW | -0.188 | ASW | -0.030 | ASW | -0.216 | ASW | AFR |
| ASW | -0.099 | ACB | -0.102 | ACB | -0.081 | ACB | -0.090 | ACB | AFR |
| LWK | 0.703 | YRI | 0.636 | YRI | 0.712 | YRI | 0.229 | YRI | AFR |
| YRI | 0.927 | LWK | 0.989 | LWK | 0.783 | LWK | 1.000 | LWK | AFR |
| CDX | 0.592 | KHV | 0.600 | KHV | 0.534 | KHV | 0.541 | KHV | EAS |
| CHB | 0.205 | CHS | 0.208 | CHS | 0.211 | CHS | 0.147 | CHS | EAS |
| CHS | 0.585 | CHB | 0.580 | CHB | 0.576 | CHB | 0.574 | CHB | EAS |
| JPT | 0.913 | CHB | 0.963 | CHB | 0.893 | CHB | 0.983 | CHB | EAS |
| KHV | 0.543 | CDX | 0.555 | CDX | 0.529 | CDX | 0.563 | CDX | EAS |
| CEU | -0.118 | GBR | -0.151 | GBR | -0.080 | GBR | -0.306 | GBR | EUR |
| FIN | 0.826 | CEU | 0.805 | CEU | 0.798 | CEU | 0.631 | TSI | EUR |
| GBR | 0.127 | CEU | 0.165 | CEU | 0.081 | CEU | 0.375 | CEU | EUR |
| IBS | 0.282 | TSI | 0.071 | TSI | 0.381 | TSI | -0.383 | TSI | EUR |
| TSI | 0.560 | IBS | 0.596 | IBS | 0.514 | IBS | 0.346 | CEU | EUR |
| GIH | 0.873 | TSI | 0.847 | FIN | 0.850 | CEU | 0.834 | CHS | SAS |

Table S13: **Region-by-region mean silhouette for the HGDP data set.** These represent averages over each population label on the space of estimated admixture coefficients, $\boldsymbol{Q}$, the higher, the better.

| | HaploADMIXTURE | | OpenADMIXTURE | | SCOPE | | TeraStructure | | |
|---|---|---|---|---|---|---|---|---|---|
| Population | Silhouette | Closest | Silhouette | Closest | Silhouette | Closest | Silhouette | Closest | Continent |
| BantuKenya | -0.341 | Mandenka | -0.347 | Mandenka | 0.492 | Yoruba | -0.351 | Mandenka | AFR |
| BantuSouthAfrica | -0.197 | Yoruba | -0.388 | Yoruba | 0.114 | BantuKenya | -0.510 | San | AFR |
| BiakaPygmy | 0.217 | MbutiPygmy | 0.000 | MbutiPygmy | 0.932 | San | 0.000 | MbutiPygmy | AFR |
| Mandenka | 0.417 | Yoruba | 0.486 | Yoruba | 0.533 | Yoruba | 0.486 | Yoruba | AFR |
| MbutiPygmy | 0.285 | BiakaPygmy | 0.000 | BiakaPygmy | 0.810 | San | 0.000 | BiakaPygmy | AFR |
| San | 0.745 | BiakaPygmy | 0.732 | MbutiPygmy | 0.894 | MbutiPygmy | -0.214 | BiakaPygmy | AFR |
| Yoruba | 0.393 | BantuSouthAfrica | 0.460 | BantuSouthAfrica | 0.731 | Mandenka | -0.105 | San | AFR |
| Colombian | 0.058 | Pima | -0.479 | Karitiana | 0.197 | Pima | -0.876 | Surui | AMR |
| Karitiana | -0.497 | Surui | -0.929 | Surui | -0.212 | Surui | -0.929 | Surui | AMR |
| Maya | 0.009 | Pima | 0.044 | Pima | -0.128 | Pima | 0.012 | Pima | AMR |
| Pima | 0.155 | Colombian | 0.126 | Colombian | 0.369 | Colombian | -0.116 | Karitiana | AMR |
| Surui | 0.476 | Karitiana | 1.000 | Karitiana | 0.097 | Karitiana | 1.000 | Karitiana | AMR |
| Balochi | -0.171 | Brahui | -0.200 | Brahui | -0.224 | Brahui | -0.190 | Brahui | CSA |
| Brahui | 0.106 | Balochi | 0.149 | Balochi | 0.136 | Balochi | 0.112 | Balochi | CSA |
| Burusho | 0.683 | Pathan | 0.632 | Pathan | 0.664 | Pathan | 0.612 | Pathan | CSA |
| Hazara | 0.099 | Uygur | 0.101 | Uygur | 0.106 | Uygur | 0.104 | Uygur | CSA |
| Kalash | 0.831 | Sindhi | 0.864 | Sindhi | 0.805 | Sindhi | 0.857 | Sindhi | CSA |
| Makrani | -0.035 | Sindhi | 0.002 | Sindhi | -0.014 | Balochi | 0.036 | Brahui | CSA |
| Pathan | 0.433 | Sindhi | 0.388 | Burusho | 0.306 | Balochi | 0.342 | Burusho | CSA |
| Sindhi | -0.019 | Brahui | 0.082 | Brahui | 0.079 | Brahui | 0.078 | Balochi | CSA |
| Uygur | 0.000 | Hazara | 0.022 | Hazara | -0.009 | Hazara | 0.012 | Hazara | CSA |
| Cambodian | 0.168 | Tu | 0.120 | Tu | 0.144 | Tu | 0.101 | Tu | EAS |
| Dai | 0.345 | Lahu | 0.574 | Lahu | 0.319 | Lahu | 0.074 | Lahu | EAS |
| Daur | 0.188 | Hezhen | 0.151 | Hezhen | 0.115 | Hezhen | 0.138 | Hezhen | EAS |
| Han | -0.295 | Tujia | -0.551 | She | -0.208 | Tujia | -0.701 | Tujia | EAS |
| Han-NChina | 0.047 | Japanese | -0.026 | Japanese | -0.004 | Japanese | -0.530 | Yi | EAS |
| Hezhen | 0.061 | Daur | 0.068 | Daur | -0.115 | Oroqen | 0.032 | Daur | EAS |
| Japanese | 0.445 | Tujia | 0.224 | Tujia | 0.164 | Tujia | -0.165 | Miao | EAS |
| Lahu | 0.205 | Dai | 0.094 | Dai | -0.064 | Dai | -0.061 | Dai | EAS |
| Miao | -0.110 | Tujia | -0.375 | Tujia | -0.077 | Tujia | -0.543 | Tujia | EAS |
| Mongola | -0.511 | Daur | -0.495 | Daur | -0.473 | Daur | -0.482 | Daur | EAS |
| Naxi | 0.175 | Yi | -0.052 | Yi | -0.042 | Yi | -0.429 | Miao | EAS |
| Oroqen | -0.058 | Daur | -0.068 | Daur | 0.053 | Hezhen | -0.171 | Daur | EAS |
| She | 0.061 | Miao | 0.381 | Tujia | 0.041 | Miao | 0.834 | Tujia | EAS |
| Tu | 0.301 | Daur | 0.274 | Daur | 0.341 | Daur | 0.240 | Daur | EAS |
| Tujia | 0.029 | Miao | -0.357 | She | -0.039 | She | -0.530 | She | EAS |
| Xibo | -0.507 | Yakut | -0.505 | Yakut | -0.446 | Yakut | -0.622 | Yakut | EAS |

Table S13: **Region-by-region mean silhouette for the HGDP data set.** These represent averages over each population label on the space of estimated admixture coefficients, $\boldsymbol{Q}$, the higher, the better.

| | HaploADMIXTURE | | OpenADMIXTURE | | SCOPE | | TeraStructure | | |
|---|---|---|---|---|---|---|---|---|---|
| Population | Silhouette | Closest | Silhouette | Closest | Silhouette | Closest | Silhouette | Closest | Continent |
| Yakut | 0.380 | Mongola | 0.373 | Mongola | 0.390 | Mongola | 0.324 | Mongola | EAS |
| Yi | -0.145 | Naxi | -0.181 | Dai | -0.161 | Dai | -0.273 | Naxi | EAS |
| Adygei | 0.747 | Tuscan | 0.766 | Tuscan | 0.717 | Russian | 0.756 | Tuscan | EUR |
| Basque | 0.782 | French | 0.787 | French | 0.717 | Italian | 0.807 | French | EUR |
| French | 0.334 | Orcadian | 0.332 | Orcadian | 0.060 | Orcadian | 0.314 | Orcadian | EUR |
| Italian | 0.540 | Tuscan | 0.517 | Tuscan | 0.216 | French | 0.526 | Tuscan | EUR |
| Orcadian | 0.733 | French | 0.725 | French | 0.496 | French | 0.671 | Basque | EUR |
| Russian | 0.717 | Orcadian | 0.738 | Orcadian | 0.686 | Orcadian | 0.641 | Orcadian | EUR |
| Sardinian | 0.829 | Italian | 0.867 | Italian | 0.753 | Basque | 0.653 | Tuscan | EUR |
| Tuscan | 0.560 | Italian | 0.539 | Italian | 0.089 | Italian | 0.412 | Sardinian | EUR |
| Bedouin | -0.252 | Palestinian | -0.252 | Palestinian | -0.133 | Palestinian | -0.181 | Palestinian | ME |
| Druze | 0.338 | Palestinian | 0.293 | Palestinian | 0.708 | Palestinian | 0.122 | Palestinian | ME |
| Mozabite | 0.494 | Palestinian | 0.535 | Palestinian | 0.528 | Bedouin | 0.400 | Palestinian | ME |
| Palestinian | 0.351 | Druze | 0.286 | Druze | 0.445 | Druze | 0.176 | Druze | ME |
| Melanesian | 0.899 | Papuan | 0.914 | Papuan | 0.901 | Papuan | 0.506 | Papuan | OCE |
| Papuan | 0.823 | Melanesian | 0.832 | Melanesian | 0.823 | Melanesian | 0.860 | Melanesian | OCE |

Table S14: **Region-by-region mean silhouette for the HO data set.** These represent averages over each population label on the space of estimated admixture coefficients, $\boldsymbol{Q}$, the higher, the better.

| | HaploADMIXTURE | | OpenADMIXTURE | | SCOPE | | TeraStructure | |
|---|---|---|---|---|---|---|---|---|
| Population | Silhouette | Closest | Silhouette | Closest | Silhouette | Closest | Silhouette | Closest |
| Abkhasian | -0.091 | Georgian | -0.113 | Georgian | -0.004 | Georgian | -0.116 | Georgian |
| Adygei | -0.195 | Balkar | -0.130 | Balkar | -0.144 | Chechen | -0.252 | Balkar |
| Albanian | 0.111 | Tuscan | -0.011 | Bulgarian | 0.038 | Bulgarian | 0.057 | Tuscan |
| Aleut | -0.374 | Mordovian | -0.366 | Mordovian | -0.359 | Russian | -0.375 | Mordovian |
| Algerian | 0.202 | Mozabite | 0.194 | Mozabite | 0.056 | Mozabite | 0.306 | Tunisian |
| Altaian | 0.460 | Tuvinian | 0.410 | Tuvinian | 0.420 | Kalmyk | 0.383 | Tuvinian |
| Ami | -0.513 | Atayal | -0.341 | Atayal | 0.170 | Atayal | -0.331 | Atayal |
| Armenian | 0.217 | Georgian Jew | 0.205 | Georgian Jew | -0.076 | Georgian Jew | -0.110 | Georgian Jew |
| Ashkenazi Jew | -0.179 | Italian South | -0.156 | Italian South | -0.182 | Italian South | -0.303 | Maltese |
| Atayal | 0.703 | Ami | 0.743 | Ami | 0.519 | Ami | 0.549 | Ami |
| Australian | 0.707 | Bougainville | 0.860 | Bougainville | 0.826 | Bougainville | 0.714 | Bougainville |
| Balkar | 0.092 | North Ossetian | 0.083 | North Ossetian | 0.021 | North Ossetian | 0.047 | North Ossetian |
| Balochi | -0.263 | Brahui | -0.207 | Brahui | -0.160 | Brahui | -0.129 | Brahui |
| BantuKenya | -0.027 | Luo | -0.042 | Luo | -0.046 | Luo | -0.413 | Ju hoan North |
| BantuSA | -0.097 | Luhya | -0.112 | Luhya | -0.095 | Luhya | -0.531 | Biaka |
| Basque | 0.133 | Spanish North | 0.264 | Spanish North | 0.082 | Spanish North | 0.034 | French South |
| BedouinA | -0.052 | Egyptian | -0.041 | Egyptian | -0.187 | Palestinian | -0.106 | Egyptian |
| BedouinB | 0.586 | Saudi | 0.566 | Saudi | 0.502 | Yemenite Jew | 0.814 | Saudi |
| Belarusian | 0.022 | Norwegian | 0.067 | Norwegian | -0.023 | Icelandic | -0.105 | Norwegian |
| Bengali | 0.691 | Punjabi | 0.520 | Punjabi | 0.436 | Punjabi | 0.715 | Punjabi |
| Bergamo | 0.519 | Bulgarian | 0.440 | Bulgarian | 0.080 | Spanish | 0.355 | Bulgarian |
| Biaka | 0.885 | Hadza | 0.887 | Hadza | 0.887 | Hadza | 0.000 | Esan |
| Bolivian | -0.265 | Quechua | -0.285 | Quechua | -0.245 | Quechua | -0.507 | Quechua |
| Bougainville | 0.841 | Australian | 0.848 | Australian | 0.735 | Australian | 0.842 | Australian |
| Brahui | 0.206 | Balochi | 0.161 | Balochi | 0.105 | Makrani | 0.056 | Balochi |
| Bulgarian | 0.277 | Bergamo | 0.327 | Croatian | 0.298 | Albanian | 0.103 | Croatian |
| Burusho | 0.501 | GujaratiA | 0.489 | GujaratiA | 0.396 | GujaratiA | 0.510 | Pathan |
| Cambodian | 0.226 | Thai | 0.205 | Thai | 0.065 | Thai | 0.190 | Thai |
| Canary Islanders | 0.437 | Sardinian | 0.245 | Sardinian | 0.476 | Spanish | 0.243 | Spanish |
| Chechen | 0.263 | Balkar | -0.030 | Lezgin | -0.033 | Lezgin | 0.006 | Adygei |
| Chukchi | 0.403 | Eskimo | 0.277 | Eskimo | 0.333 | Eskimo | 0.424 | Eskimo |
| Chuvash | 0.552 | Saami WGA | 0.616 | Saami WGA | 0.541 | Saami WGA | 0.417 | Saami WGA |
| Cochin Jew | -0.438 | GujaratiB | -0.342 | GujaratiB | -0.532 | GujaratiB | -0.359 | GujaratiB |
| Croatian | 0.472 | Hungarian | 0.441 | Hungarian | 0.303 | Hungarian | 0.085 | Hungarian |
| Cypriot | 0.140 | Lebanese | 0.069 | Lebanese | -0.024 | Turkish Jew | 0.122 | Turkish Jew |
| Czech | 0.008 | Scottish | -0.060 | Orcadian | -0.184 | Scottish | -0.104 | Orcadian |
| Dai | 0.225 | Ami | 0.200 | Kinh | 0.218 | Kinh | -0.128 | Ami |

Table S14: **Region-by-region mean silhouette for the HO data set.** These represent averages over each population label on the space of estimated admixture coefficients, $\boldsymbol{Q}$, the higher, the better.

| | HaploADMIXTURE | | OpenADMIXTURE | | SCOPE | | TeraStructure | |
|---|---|---|---|---|---|---|---|---|
| Population | Silhouette | Closest | Silhouette | Closest | Silhouette | Closest | Silhouette | Closest |
| Datog | 0.721 | Somali | 0.742 | Somali | 0.669 | Somali | 0.830 | Masai |
| Daur | 0.686 | Hezhen | 0.607 | Hezhen | 0.525 | Hezhen | 0.552 | Hezhen |
| Dolgan | 0.036 | Yakut | -0.043 | Yakut | -0.123 | Yakut | -0.016 | Yakut |
| Druze | -0.100 | Iraqi Jew | 0.015 | Iraqi Jew | 0.312 | Iraqi Jew | 0.376 | Iranian Jew |
| Egyptian | 0.335 | Yemenite Jew | 0.387 | Yemenite Jew | 0.368 | BedouinA | 0.230 | Yemenite Jew |
| English | 0.004 | Orcadian | 0.040 | Orcadian | 0.110 | Czech | -0.154 | Orcadian |
| Esan | -0.117 | Yoruba | -0.141 | Yoruba | -0.025 | Yoruba | 0.000 | Biaka |
| Eskimo | 0.598 | Chukchi | 0.594 | Chukchi | 0.495 | Chukchi | 0.677 | Chukchi |
| Estonian | 0.047 | Lithuanian | -0.090 | Lithuanian | -0.101 | Lithuanian | 0.245 | Lithuanian |
| Ethiopian Jew | 0.772 | Somali | 0.750 | Somali | 0.703 | Somali | 0.859 | Somali |
| Even | -0.611 | Mansi | -0.560 | Mansi | -0.522 | Mansi | -0.362 | Mansi |
| Finnish | 0.193 | Russian | 0.113 | Estonian | 0.002 | Estonian | -0.087 | Russian |
| French | -0.091 | English | -0.035 | English | -0.095 | Croatian | -0.115 | English |
| French South | 0.010 | Basque | 0.035 | Spanish North | -0.131 | Spanish North | 0.044 | Spanish North |
| Gambian | -0.111 | Mandenka | -0.011 | Mandenka | 0.109 | Mandenka | -0.491 | Mandenka |
| Georgian | 0.250 | Abkhasian | 0.208 | Abkhasian | 0.206 | Abkhasian | 0.084 | Abkhasian |
| Georgian Jew | 0.021 | Iranian Jew | 0.091 | Iranian Jew | 0.042 | Armenian | -0.056 | Iranian Jew |
| Greek | -0.557 | Tuscan | -0.494 | Tuscan | -0.425 | Albanian | -0.436 | Tuscan |
| GujaratiA | -0.074 | GujaratiB | -0.002 | GujaratiB | 0.265 | Sindhi | -0.018 | GujaratiB |
| GujaratiB | 0.165 | GujaratiC | 0.104 | GujaratiC | 0.288 | GujaratiC | 0.228 | GujaratiC |
| GujaratiC | 0.216 | Punjabi | 0.274 | GujaratiB | 0.051 | GujaratiB | -0.077 | GujaratiB |
| GujaratiD | 0.262 | Punjabi | 0.174 | Punjabi | 0.004 | GujaratiC | 0.480 | Punjabi |
| Hadza | 0.782 | Biaka | 0.815 | Biaka | 0.829 | Biaka | 0.782 | Kikuyu |
| Han | -0.638 | Miao | -0.595 | She | -0.441 | She | -0.586 | Miao |
| Han NChina | -0.001 | Naxi | -0.032 | Naxi | 0.060 | Naxi | -0.255 | Naxi |
| Hazara | 0.368 | Uygur | 0.342 | Uygur | 0.348 | Uygur | 0.166 | Uygur |
| Hezhen | -0.441 | Xibo | -0.427 | Daur | -0.422 | Xibo | -0.366 | Xibo |
| Hungarian | 0.006 | French | -0.043 | Croatian | -0.090 | Croatian | -0.085 | Croatian |
| Icelandic | -0.271 | Belarusian | -0.178 | Belarusian | -0.084 | Norwegian | -0.211 | Scottish |
| Iranian | 0.570 | Georgian Jew | 0.511 | Georgian Jew | 0.258 | Georgian | 0.336 | Georgian Jew |
| Iranian Jew | -0.033 | Iraqi Jew | -0.046 | Iraqi Jew | -0.118 | Iraqi Jew | 0.192 | Iraqi Jew |
| Iraqi Jew | 0.349 | Iranian Jew | 0.352 | Iranian Jew | 0.335 | Iranian Jew | -0.085 | Georgian Jew |
| Italian South | 0.000 | Ashkenazi Jew | 0.000 | Sicilian | 0.000 | Ashkenazi Jew | 0.000 | Sicilian |
| Itelmen | 0.581 | Koryak | 0.577 | Koryak | 0.415 | Koryak | -0.075 | Koryak |
| Japanese | 0.199 | Korean | 0.148 | Korean | 0.096 | Korean | 0.256 | Korean |
| Jordanian | -0.007 | Palestinian | -0.048 | Palestinian | 0.042 | Palestinian | -0.012 | Palestinian |
| Ju hoan North | 0.999 | Khomani | 1.000 | Khomani | 0.922 | Khomani | 0.381 | BantuSA |

Table S14: **Region-by-region mean silhouette for the HO data set.** These represent averages over each population label on the space of estimated admixture coefficients, $\boldsymbol{Q}$, the higher, the better.

| | HaploADMIXTURE | | OpenADMIXTURE | | SCOPE | | TeraStructure | |
|---|---|---|---|---|---|---|---|---|
| Population | Silhouette | Closest | Silhouette | Closest | Silhouette | Closest | Silhouette | Closest |
| Kalash | 0.760 | GujaratiC | 0.673 | GujaratiD | 0.581 | GujaratiA | 0.936 | Turkmen |
| Kalmyk | 0.607 | Kyrgyz | 0.558 | Altaian | 0.446 | Altaian | 0.500 | Kyrgyz |
| Karitiana | 1.000 | Surui | 1.000 | Surui | 0.931 | Piapoco | 0.000 | Piapoco |
| Khomani | -0.442 | Mbuti | -0.424 | Mbuti | -0.274 | Mbuti | -0.550 | Luhya |
| Kikuyu | 0.658 | Masai | 0.642 | Masai | 0.548 | Masai | 0.636 | Hadza |
| Kinh | 0.065 | Dai | 0.085 | Dai | 0.038 | Lahu | -0.264 | Dai |
| Korean | -0.222 | Naxi | -0.105 | Han NChina | -0.033 | Han NChina | 0.032 | Naxi |
| Koryak | -0.344 | Itelmen | -0.328 | Itelmen | -0.223 | Itelmen | 0.020 | Itelmen |
| Kumyk | -0.198 | Lezgin | -0.059 | Lezgin | -0.153 | North Ossetian | 0.030 | Lezgin |
| Kusunda | 0.607 | Uygur | 0.626 | Uygur | 0.615 | Hazara | 0.460 | Thai |
| Kyrgyz | 0.597 | Kalmyk | 0.523 | Altaian | 0.449 | Altaian | 0.524 | Kalmyk |
| LaBrana | 0.000 | Russian | 0.000 | Lithuanian | 0.000 | Lithuanian | 0.000 | Lithuanian |
| Lahu | -0.004 | She | 0.115 | She | 0.129 | She | -0.189 | She |
| Lebanese | -0.201 | Palestinian | -0.222 | Palestinian | -0.348 | BedouinA | -0.207 | Palestinian |
| Lezgin | 0.207 | Chechen | 0.284 | Chechen | 0.085 | Chechen | -0.025 | Kumyk |
| Libyan Jew | 0.285 | Tunisian Jew | 0.311 | Tunisian Jew | 0.109 | Tunisian Jew | 0.067 | Tunisian Jew |
| Lithuanian | 0.295 | Estonian | 0.099 | Estonian | 0.108 | Estonian | 0.340 | Estonian |
| Luhya | -0.165 | Luo | -0.160 | Luo | -0.090 | Luo | 0.002 | Luo |
| Luo | 0.129 | Luhya | 0.040 | Luhya | 0.052 | Luhya | 0.355 | BantuKenya |
| Makrani | 0.302 | Brahui | 0.227 | Brahui | 0.215 | Brahui | 0.099 | Brahui |
| Maltese | -0.071 | Sicilian | -0.028 | Sicilian | -0.239 | Italian South | 0.053 | Sicilian |
| Mandenka | 0.474 | Esan | 0.466 | Esan | 0.095 | Yoruba | -0.005 | BantuSA |
| Mansi | 0.551 | Tubalar | 0.567 | Tubalar | 0.584 | Tubalar | 0.546 | Selkup |
| Masai | 0.583 | Kikuyu | 0.531 | Kikuyu | 0.436 | Kikuyu | 0.594 | Kikuyu |
| Mayan | 0.026 | Zapotec | 0.031 | Zapotec | 0.005 | Zapotec | -0.397 | Zapotec |
| Mbuti | 0.834 | Khomani | 0.859 | Khomani | 0.868 | Khomani | 0.000 | Biaka |
| Mende | 0.710 | Gambian | 0.686 | Gambian | 0.460 | Gambian | -0.904 | Biaka |
| Miao | 0.374 | She | 0.158 | She | 0.020 | She | 0.187 | She |
| Mixe | 0.444 | Mixtec | 0.565 | Mixtec | 0.381 | Mixtec | -0.580 | Karitiana |
| Mixtec | -0.417 | Mixe | -0.242 | Mixe | 0.002 | Mixe | -0.160 | Pima |
| Mongola | 0.060 | Xibo | 0.070 | Xibo | -0.027 | Xibo | -0.044 | Xibo |
| Mordovian | 0.293 | Russian | 0.216 | Russian | 0.195 | Russian | 0.024 | Russian |
| Moroccan Jew | 0.456 | Turkish Jew | 0.482 | Turkish Jew | 0.270 | Turkish Jew | 0.031 | Turkish Jew |
| Mozabite | -0.068 | Saharawi | -0.033 | Saharawi | -0.071 | Saharawi | -0.075 | Saharawi |
| Naxi | 0.139 | Han NChina | 0.146 | Han NChina | 0.152 | Yi | 0.042 | Han NChina |
| Nganasan | 0.984 | Dolgan | 0.899 | Dolgan | 0.851 | Dolgan | 0.850 | Saqqaq |
| Nogai | 0.409 | Kumyk | 0.451 | Turkmen | 0.365 | Balkar | 0.382 | Kumyk |

Table S14: **Region-by-region mean silhouette for the HO data set.** These represent averages over each population label on the space of estimated admixture coefficients, $\boldsymbol{Q}$, the higher, the better.

| | HaploADMIXTURE | | OpenADMIXTURE | | SCOPE | | TeraStructure | |
|---|---|---|---|---|---|---|---|---|
| Population | Silhouette | Closest | Silhouette | Closest | Silhouette | Closest | Silhouette | Closest |
| North Ossetian | -0.039 | Balkar | -0.073 | Balkar | -0.019 | Balkar | -0.125 | Chechen |
| Norwegian | 0.117 | Orcadian | 0.017 | Scottish | -0.065 | Scottish | -0.081 | Icelandic |
| Orcadian | -0.019 | Norwegian | -0.017 | Scottish | -0.072 | Scottish | 0.056 | Czech |
| Oroqen | -0.431 | Yakut | -0.428 | Yakut | -0.397 | Yakut | -0.411 | Ulchi |
| Palestinian | 0.130 | Jordanian | 0.097 | Jordanian | 0.090 | Jordanian | 0.043 | Jordanian |
| Papuan | 1.000 | Australian | 1.000 | Australian | 0.936 | Australian | 1.000 | Australian |
| Pathan | 0.331 | Burusho | 0.297 | Burusho | 0.265 | GujaratiA | 0.036 | Balochi |
| Piapoco | 0.903 | Bolivian | 0.910 | Bolivian | 0.749 | Bolivian | 0.000 | Karitiana |
| Pima | 0.910 | Mixe | 0.829 | Mixe | 0.701 | Mixe | -0.563 | Mixtec |
| Punjabi | -0.229 | GujaratiD | -0.218 | GujaratiD | -0.142 | GujaratiD | -0.211 | GujaratiD |
| Quechua | -0.017 | Mayan | -0.023 | Mayan | 0.057 | Bolivian | -0.201 | Pima |
| Russian | 0.134 | Mordovian | 0.102 | Mordovian | 0.007 | Mordovian | 0.019 | Mordovian |
| Saami WGA | 0.000 | Chuvash | 0.000 | Russian | 0.000 | Chuvash | 0.000 | Chuvash |
| Saharawi | 0.521 | Mozabite | 0.523 | Mozabite | 0.404 | Mozabite | 0.386 | Mozabite |
| Saqqaq | 0.000 | Eskimo | 0.000 | Koryak | 0.000 | Koryak | 0.000 | Eskimo |
| Sardinian | 0.486 | Canary Islanders | 0.647 | Canary Islanders | 0.780 | Canary Islanders | 0.522 | Tuscan |
| Saudi | -0.289 | BedouinB | -0.313 | BedouinB | -0.529 | BedouinB | -0.527 | BedouinB |
| Scottish | -0.383 | Orcadian | -0.278 | Orcadian | -0.117 | Orcadian | -0.352 | Orcadian |
| Selkup | -0.222 | Mansi | -0.198 | Mansi | -0.191 | Mansi | -0.119 | Mansi |
| She | 0.034 | Dai | 0.148 | Kinh | 0.056 | Kinh | 0.259 | Dai |
| Sicilian | -0.011 | Maltese | -0.057 | Maltese | -0.278 | Italian South | -0.166 | Maltese |
| Sindhi | 0.158 | Burusho | 0.110 | GujaratiA | 0.187 | GujaratiB | 0.017 | GujaratiB |
| Somali | 0.674 | Datog | 0.665 | Datog | 0.592 | Datog | 0.776 | Datog |
| Spanish | 0.255 | Canary Islanders | 0.238 | Canary Islanders | 0.186 | Bergamo | 0.089 | Canary Islanders |
| Spanish North | 0.291 | French South | -0.058 | French South | 0.178 | French South | 0.065 | French South |
| Surui | -0.875 | Karitiana | -0.570 | Karitiana | 0.887 | Piapoco | 0.000 | Karitiana |
| Syrian | -0.273 | Jordanian | -0.203 | Jordanian | -0.219 | Jordanian | -0.196 | Jordanian |
| Tajik Pomiri | 0.616 | Turkmen | 0.574 | Turkmen | 0.477 | Lezgin | 0.482 | Turkmen |
| Thai | -0.054 | Cambodian | -0.055 | Cambodian | 0.045 | Cambodian | -0.098 | Cambodian |
| Tlingit | 0.175 | Aleut | 0.191 | Aleut | 0.163 | Aleut | 0.158 | Aleut |
| Tu | 0.390 | Japanese | 0.363 | Japanese | 0.300 | Naxi | 0.213 | Japanese |
| Tubalar | 0.689 | Altaian | 0.661 | Altaian | 0.619 | Altaian | 0.749 | Selkup |
| Tujia | 0.011 | Miao | 0.009 | Miao | 0.037 | Miao | -0.291 | Miao |
| Tunisian | 0.301 | Algerian | 0.309 | Algerian | 0.082 | Egyptian | -0.124 | Algerian |
| Tunisian Jew | -0.127 | Libyan Jew | -0.197 | Libyan Jew | -0.104 | Libyan Jew | -0.084 | Libyan Jew |
| Turkish | -0.297 | Cypriot | -0.206 | Cypriot | -0.198 | Armenian | -0.119 | Georgian Jew |
| Turkish Jew | 0.085 | Moroccan Jew | 0.148 | Moroccan Jew | -0.000 | Moroccan Jew | 0.136 | Moroccan Jew |

Table S14: **Region-by-region mean silhouette for the HO data set.** These represent averages over each population label on the space of estimated admixture coefficients, $\boldsymbol{Q}$, the higher, the better.

| | HaploADMIXTURE | | OpenADMIXTURE | | SCOPE | | TeraStructure | |
|---|---|---|---|---|---|---|---|---|
| Population | Silhouette | Closest | Silhouette | Closest | Silhouette | Closest | Silhouette | Closest |
| Turkmen | 0.154 | Uzbek | 0.127 | Uzbek | -0.017 | Uzbek | 0.091 | Uzbek |
| Tuscan | 0.491 | Albanian | 0.379 | Albanian | 0.100 | Albanian | 0.047 | Sardinian |
| Tuvinian | 0.295 | Altaian | 0.281 | Altaian | 0.325 | Kalmyk | -0.161 | Altaian |
| Ukrainian | -0.137 | Belarusian | -0.058 | Belarusian | -0.112 | Belarusian | 0.077 | Scottish |
| Ulchi | 0.402 | Tuvinian | 0.436 | Tuvinian | 0.468 | Tuvinian | 0.648 | Oroqen |
| Uygur | -0.011 | Hazara | -0.048 | Hazara | -0.067 | Hazara | 0.007 | Hazara |
| Uzbek | -0.015 | Hazara | -0.061 | Hazara | -0.128 | Hazara | -0.147 | Turkmen |
| Xibo | 0.482 | Mongola | 0.344 | Mongola | 0.291 | Mongola | 0.150 | Mongola |
| Yakut | 0.476 | Dolgan | 0.431 | Dolgan | 0.434 | Dolgan | 0.448 | Tuvinian |
| Yemen | -0.310 | Egyptian | -0.336 | Egyptian | -0.429 | Egyptian | -0.280 | Egyptian |
| Yemenite Jew | 0.662 | BedouinA | 0.592 | Saudi | 0.490 | Saudi | 0.520 | BedouinA |
| Yi | -0.439 | Naxi | -0.430 | Naxi | -0.316 | Naxi | -0.470 | Naxi |
| Yoruba | 0.204 | Esan | 0.232 | Esan | -0.013 | Esan | -0.957 | Biaka |
| Yukagir | -0.436 | Dolgan | -0.485 | Dolgan | -0.412 | Dolgan | -0.568 | Nganasan |
| Zapotec | 0.162 | Mixe | 0.186 | Mixtec | 0.011 | Mixtec | 0.205 | Mixtec |

Table S15: **Root-mean-square error of SKFR from the baseline for HaploADMIXTURE and OpenADMIXTURE on the HGDP data set.** Root-mean-square error (RMSE) from baseline compares estimated admixture coefficients of sparse $K$-means (SKFR) to those estimated using all the SNPs; the lower, the better.

| SNPs | HaploADMIXTURE | OpenADMIXTURE |
|---|---|---|
| 100,000 | 0.076 | 0.172 |
| 80,000 | 0.060 | 0.173 |
| 60,000 | 0.040 | 0.174 |
| 40,000 | 0.042 | 0.179 |
| 20,000 | 0.066 | 0.177 |
| 10,000 | 0.074 | 0.180 |
| 5000 | 0.080 | 0.081 |

Table S16: **Root-mean-square error of SKFR from the baseline for HaploADMIXTURE and OpenADMIXTURE on the HO data set.** Root-mean-square error (RMSE) from baseline compares estimated admixture coefficients of sparse $K$-means (SKFR) to those estimated using all the SNPs; the lower, the better.

| SNPs | HaploADMIXTURE | OpenADMIXTURE |
|---|---|---|
| 100,000 | 0.015 | 0.021 |
| 80,000 | 0.014 | 0.024 |
| 60,000 | 0.018 | 0.027 |
| 40,000 | 0.022 | 0.034 |
| 20,000 | 0.086 | 0.093 |
| 10,000 | 0.093 | 0.126 |
| 5000 | 0.108 | 0.150 |

Table S17: **Average software runtime on each data set.** Time is in hour:minute. Sparse $K$-means (SKFR) is not applied.

| Software | TGP | HGDP | HO |
|---|---|---|---|
| HaploADMIXTURE | 2:08 | 0:31 | 1:08 |
| OpenADMIXTURE | 0:19 | 0:02 | 0:08 |
| SCOPE | 0:10 | 0:01 | 0:02 |
| TeraStructure | 1:13 | 0:09 | 0:39 |
